# Aberrant influenza virus transcription termination generates capped cRNAs that activate RIG-I

**DOI:** 10.1101/2024.11.12.623191

**Authors:** Elizaveta Elshina, Michael S. Oade, Emmanuelle Pitre, Marisa Mendes, Ahn Young Jeong, Jingru Fang, Karishma Bisht, Brandon Schweibenz, Rebecca L.Y. Fan, Hollie French, Ji Woo Park, Wei Wang, Leo L.M. Poon, Joseph Marcotrigiano, Alistair B. Russell, Aartjan J.W. te Velthuis

**Author notes:** School of Biodiversity, One Health and Veterinary Medicine, University of Glasgow, Glasgow, G12 8QQ, United Kingdom (Hollie French).

## Abstract

Host pathogen receptor retinoic acid-inducible gene I (RIG-I) detects influenza virus genome segments and non-canonical replication products. However, it has also been suggested that viral transcription products trigger innate immune responses. In this study, we investigated whether a non-canonical viral transcript is produced during infection and under what conditions this RNA is immunostimulatory. We show that for a minority of transcripts, the wild-type viral polymerase reads through the polyadenylation signal generating a capped complementary RNA (ccRNA) that contains a 5ʹ cap and the 3ʹ terminus of a cRNA. These ccRNAs are produced *in vitro*, in ribonucleoprotein reconstitution assays, in cell culture, and in murine infection models. Non-canonical transcription termination is enhanced by mutations in the triple stranded β-sheet of the viral polymerase PB1 subunit. We further show that ccRNA production correlates with RIG-I-dependent innate immune activation and that ccRNAs are bound by RIG-I in infection. Notably, while ccRNAs are not inherently immunostimulatory, they can form a RIG-I agonist by hybridizing with a complementary negative-sense viral RNA. These findings thus identify a mechanism by which IAV transcriptional products contribute to innate immune activation during infection.

## Introduction

Innate immune signalling plays a pivotal role in determining the outcome of influenza A virus (IAV) infection and can either trigger a protective or dysregulated antiviral response^1,2^. While it is established that retinoic acid-inducible gene I (RIG-I) is crucial for the detection of IAV infection^3–6^, the mechanisms underlying this recognition remain incompletely understood. Optimal RIG-I ligands are known to be double-stranded RNAs (dsRNA) with a 5ʹ tri- or diphosphate^7–9^. During IAV infection, such a structure is thought to be provided by the viral promoter, which consists of partially complementary 5ʹ and 3ʹ genome termini that contain a 5ʹ-triphosphate^10,11^. However, inside the cells viral RNA primarily exists in viral ribonucleoprotein complexes (RNP), in which the viral promoter is bound by a heterotrimeric RNA polymerase, consisting of subunits PB1, PB2 and PA, and the rest of the RNA is coated by multiple copies of the viral nucleoprotein (NP)^12,13^. It therefore remains unclear how RIG-I can access IAV RNA during an infection. Indeed, single-cell sequencing studies suggest that RIG-I activation during infection by a lab-adapted IAV strain is a rare and stochastic event even in the absence the viral immune modulator NS1^14–16^.

Active viral RNA synthesis is important for innate immune recognition during an IAV infection^17^. During replication, IAV negative-sense viral RNA (vRNA) is copied via a complementary RNA (cRNA) intermediate or transcribed into viral mRNAs. Viral replication initiation is an unprimed process that produces a full, complementary copy of each vRNA (reviewed in^18^), whereas transcription is a primed process that is initiated after cleavage of the first 8-14 nt of capped host (pre-)mRNAs^19–21^. Viral transcription termination occurs when the RNA polymerase stutters on a uridine stretch (U-stretch) ∼16-22 nt from the 5ʹ end of the vRNA template and adds a poly(A) tail to the nascent IAV mRNA^22,23^.

Viral replication can also result in the production of non-canonical RNAs such as deletion-containing viral genomes (DelVG), mini viral RNAs (mvRNA), and small viral RNAs (reviewed in^24^). Of those, DelVGs and mvRNAs, but not svRNAs, have been shown to be RIG-I agonists^25–28^. Interestingly, innate immune activation can occur under circumstances where only primary transcription can occur such as during chemical inhibition of IAV replication^29–31^ or during infection with a single-cycle IAV incapable of replication^32^. Increased IFN expression has also been observed when viral transcription is enhanced, such as after depletion of Negative Elongation Factor (NELF) complex^33^. This suggests that viral transcripts are capable of innate immune activation, even though full-length viral mRNAs are protected from innate immune recognition by their 5ʹ cap^34,35^.

To investigate how IAV transcription may contribute to innate immune signalling, we explored i) whether the IAV RNA polymerase generates non-canonical transcripts during infection and ii) whether and how such transcripts can be immunostimulatory. Our findings reveal that the viral polymerase can bypass the U-stretch during transcription termination, producing capped transcripts with a cRNA 3ʹ terminus, or capped cRNAs (ccRNA). During infection these non-canonical transcripts are detected by RIG-I along with other known viral replication products. We further show that ccRNAs can activate RIG-I when their 3ʹ terminus is hybridized to the triphosphorylated 5ʹ terminus of the negative-sense RNAs. Collectively, we suggest a new mechanism by which non-canonical viral transcription can contribute to innate immune activation during IAV infection.

## Results

### T677A mutation in PB1 induces aberrant transcription termination

The promoter proximal region of the IAV RNA polymerase undergoes significant restructuring as the enzyme transitions from a transcription pre-initiation to elongation/termination conformation, with the PB1 β-ribbon forming a triple-stranded β-sheet with PB1 residues 676-678 (Fig. 1a). To assess the role of this region on transcription termination we made six alanine substitutions in the promoter-proximal PB1 residues (i.e., R670A, N671A, S673A, N676A, T677A and S678A) and used an *in vitro* transcription assay in which the wild-type or mutant IAV RNA polymerases copied a segment 6-based 71 nt-long RNA (NA71). This RNA template had the conserved 5ʹ and 3ʹ genome termini of segment 6 and a large internal deletion that allowed resolution of the transcription products by PAGE. Suspected polyadenylated transcripts appeared as a smear above the 80 nt ssDNA marker. We also noticed two single bands at the 80 and 90 nt markers. Based on the sizes of those products we speculate that the 80 nt band likely represents a non-polyadenylated transcript that terminated at the vRNA U-stretch, while the 90 nt band likely represents a read-through transcript that terminated with a cRNA 3ʹ terminus (Fig. 1b, Extended Data Fig. 1a). (Note that the capped RNA products migrate slightly differently than the ssDNA marker due to the presence of additional phosphates, and an m^7^G cap). Of all PB1 mutations tested, T677A most potently reduced the intensity of the suspected polyadenylated transcripts, while increasing the intensity of the aberrant read-through product (Fig. 1b).

**Fig. 1.**
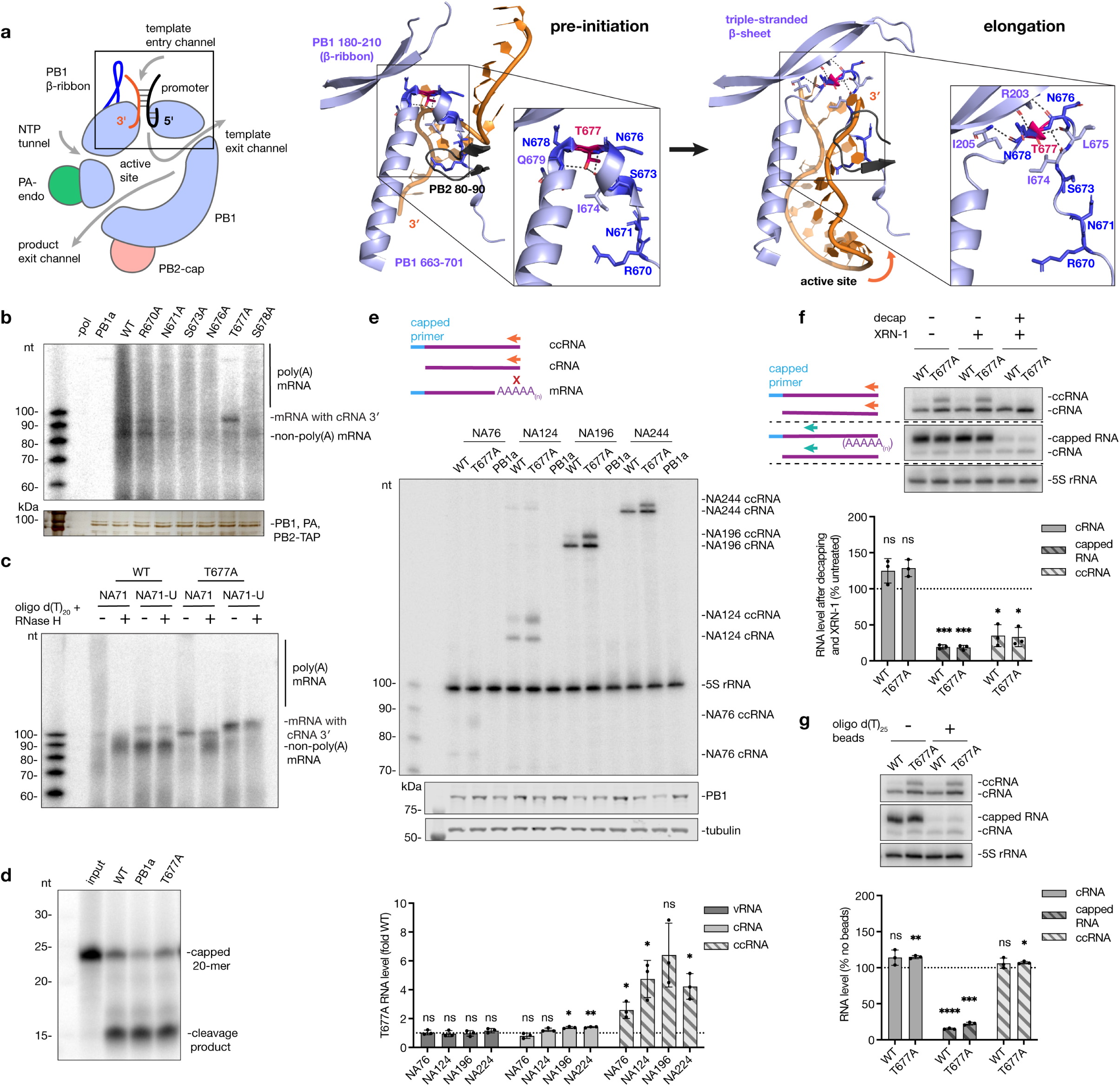
Characterization of transcription termination and ccRNA synthesis. **a**, Schematic of the IAV RNA polymerase with the promoter proximal region boxed (left); structural comparison of the promoter-proximal region in the bat IAV polymerase pre-initiation (PDB: 6T0N, center) and elongation (PDB: 6T0V, right) complexes. Mutated PB1 residues are shown in dark blue, T677 is coloured magenta, and other residues that interact with T677 or involved in the triple-stranded β-sheet formation are labelled light blue. Hydrogen bonds formed by T677 and those involved in triple-stranded β-sheet formation are indicated by black dotted lines. Note, termination conformation (PDB: 6TW1) is similar to elongation conformation and therefore not shown. **b**, *In vitro* transcription assay on a model NA71 template with purified wild-type or mutant polymerases (top). Inactive polymerase mutant (PB1a) was used as a negative control. Purified IAV polymerases visualised on a silver-stained SDS-PAGE gel (bottom). **c**, *In vitro* transcription assay with purified wild-type or T677A polymerases on a NA71 template or NA71 with a disrupted U-stretch (NA71-U), with and without oligo d(T)_20_-RNase H treatment. **d**, *In vitro* cap cleavage assay of a capped 20-mer in the presence of the model vRNA promoter. **e**, RNP reconstitution assay with short segment 6-based templates: schematic of primer extension for cRNA and ccRNA with a terminal cRNA primer (NA 5′) (orange, top; ‘x’ indicates no binding to the mRNA); primer extension gel and western blot analysis of PB1 expression levels (middle); quantification of vRNA, cRNA and ccRNA levels produced by the T677A mutant versus wild-type polymerase (bottom). Primer extension schematic and a representative gel for vRNA is provided in Supplementary Fig. 1a, b. **f**, Enzymatic digestion of total RNA from RNP reconstitution with the NA196 template: schematic of primer extension with terminal (orange) and internal (green) primers (top left); primer extension gel (top right); quantification of cRNA, all capped RNA, and ccRNA levels post-decapping and XRN-1 treatment (bottom). **g**, Oligo d(T)_25_-bead-based depletion of polyadenylated RNAs in total RNA from an RNP reconstitution assay with the NA196 template: primer extension gel with terminal (NA 5′) and internal (NA c/mRNA mini) primers (top); quantification of cRNA, all capped RNA, and ccRNA levels post-treatment (bottom). All gel images (except silver-stained SDS-PAGE in **b**, since polymerases have been purified once) are representative of three independent replicates. Data in panels **e**-**g** are presented as mean ± SD from three independent experiments. Statistical significance was determined using a one-sample t-test where each condition was compared to a theoretical mean of 1 or 100 (grouping is for visual purposes only, no comparisons have been made between groups) (ns=non-significant, \**P*<0.05, \*\**P*<0.01, \*\*\**P*<0.001, \*\*\*\**P*<0.0001).

To further characterize the products of the *in vitro* reactions, transcription products were incubated with oligo d(T)_20_ and treated with RNase H to degrade poly(A) tails. The oligo d(T)_20_-RNase H treatment removed the slower migrating smear and increased the intensity of the suspected non-polyadenylated mRNA product (Fig. 1c). As positive control for the identity of the aberrant read-through product, we used an NA71 template with an interrupted U-stretch (NA71-U), whereby 5ʹ-UUUUUU was replaced by 5ʹ-AUGUGU. This template modification prevents polymerase stuttering and polyadenylation and instead allows the polymerase to read though the U-stretch and copy the cRNA 3ʹ terminus (Supplementary Table 9). Transcription of the NA71-U template by the wild-type polymerase resulted in accumulation of an aberrant product of a similar size to that produced by the T677A polymerase on the NA71 template (Fig. 1c). This confirmed that the aberrant product generated by the T677A polymerase is likely a U-stretch read-though transcript that is terminated with a cRNA 3ʹ end instead of a poly(A) tail (Fig. 1c, Extended Data Fig. 1a). *In vitro* activity assays on a model promoter showed that the T677A mutant was able to cleave a capped primer with equal efficiency as the wild-type polymerase (Fig. 1d), suggesting that the T677A mutation only affected transcription termination.

### Aberrant transcription termination results in capped cRNA synthesis

To test whether aberrant transcription termination can indeed generate mRNAs containing a cRNA 3ʹ terminus instead of poly(A) tail in cell culture, we used wild-type and T677A mutant polymerases in an RNP assay with short segment 6-based templates that contained internal deletions, similar to DelVGs and mvRNAs. Primer extension analysis using a primer complementary to the cRNA 3ʹ-terminal nucleotides, which are absent in mRNAs, showed two bands that migrated approximately 10-12 nt apart (Fig. 1e, Supplementary Fig. 1a). The lower band migrated at the expected size of a cRNA, and the upper band at the size of a cRNA with a 10-12 nt capped primer, which likely represents the aberrant transcription product with a cRNA 3ʹ terminus that we observed *in vitro* (Fig. 1b, c). We will call this non-canonical transcription product capped cRNA (ccRNA) from hereon. The T677A mutation significantly increased ccRNA synthesis on most short templates, but it had no significant effect on the vRNA synthesis and cRNA production was only modestly affected (Fig. 1e, Supplementary Fig. 1b). When tested with full-length templates, the T677A mutation also modestly enhanced vRNA and cRNA synthesis, but not total capped RNA synthesis (including mRNA and ccRNA) (Supplementary Fig. 1c, d).

To verify that the ccRNA contains a cap, we first used enzymatic decapping followed by XRN-1 exoribonuclease digestion to remove all capped transcripts. Next, we measured cRNA and ccRNA levels with a cRNA 3ʹ-terminus specific primer and detected total capped RNA levels with an internal primer that aligns to mRNA, ccRNA, and cRNA molecules (Fig. 1f, Supplementary Fig. 1a). Decapping-based digestion did not affect cRNA levels as they are protected from XRN-1 digestion by the 5ʹ-triphosphate group, while levels of total capped RNAs and ccRNAs were significantly reduced, confirming that ccRNAs are capped (Fig. 1f). Depletion of polyadenylated RNA with oligo d(T)_25_ beads reduced capped RNA levels by 75-85%, while cRNA and ccRNA levels were not reduced, confirming that ccRNAs are not polyadenylated (Fig. 1g).

PB1 T677 is a highly conserved residue that forms dynamic interactions in the triple-stranded β-sheet region as the polymerase transitions between transcriptional stages and its mutation thus likely affects the activity of the viral polymerase (Fig. 1a, Extended Data Fig. 1b). To investigate if ccRNA formation was alanine substitution-specific, we generated four additional amino acid substitutions. Three of these four substitutions also elevated ccRNA production on a 196 nt-long segment 6 (NA196) template (Extended Data Fig. 1c), highlighting the importance of PB1 T677 in transcription termination. To confirm that ccRNA formation is not specific to the polymerase of the A/WSN/33 strain, we performed cell-based activity assays that showed that ccRNAs were also produced by the wild-type RNA polymerases of some, but not all seasonal and pandemic IAV strains (Extended Data Fig. 1d). Taken together, the above results confirm that both wild-type and to a larger degree the T677A mutant polymerase can generate non-canonical transcripts containing a 5ʹ cap and a 3ʹ cRNA terminus.

### ccRNAs are produced in viral infection

As primer extension analyses can only provide nucleotide resolution for shorter templates, we employed a template switching oligonucleotide (TSO)-based RT-PCR, which uses a reverse transcriptase to append a known sequence to the 5ʹ termini of cDNA molecules, to determine whether ccRNAs are produced during infection. First, cRNA and ccRNA were reverse-transcribed using cRNA 3ʹ-terminus specific primer and then amplified using a TSO-specific forward and segment-specific reverse primers. The products were subsequently separated by PAGE into faster migrating cRNA and slower migrating ccRNA molecules (Fig. 2a). We first tested this method using various *in vitro* transcribed RNAs that were derived from a 196 nt long segment 6-based template (NA196), which, like NA71, contains 5ʹ and 3ʹ genome termini but a slightly shorter internal deletion. We found that while the method could successfully distinguish between cRNAs and ccRNAs, it showed a weak, unspecific amplification of capped RNAs that lacked the RT primer binding site. The ccRNA specificity was improved in reactions where both cRNA and capped RNAs were present (which is more representative of infection), but unspecific amplification was still observed, likely due to reverse transcriptase switching between identical internal sequences (Extended Data Fig. 2a, b). We therefore used the TSO assay only for the initial identification of ccRNAs and implemented a poly(A) RNA depletion step to minimize unspecific amplification (see below).

**Fig. 2.**
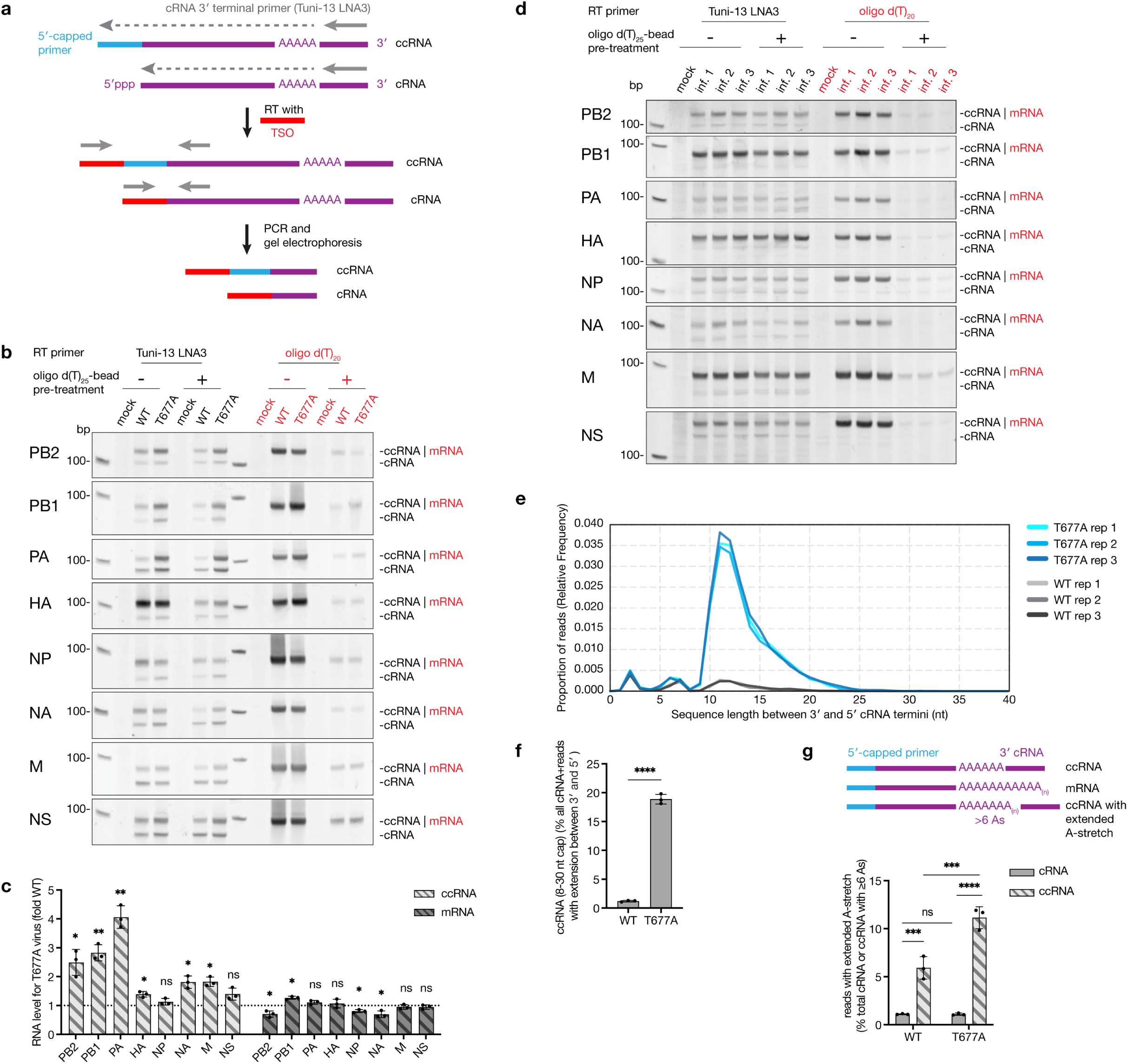
Detection of ccRNA produced during IAV infection. **a,** Schematic of the TSO-based RT-PCR for analysis of 5ʹ cRNA termini. RT was done with a universal primer containing LNA modifications (Tuni-13 LNA3), which binds to the conserved 3ʹ cRNA termini of all eight segments, and in the presence of TSO. The TSO was appended by the RT enzyme to the 5ʹ termini of cDNA molecules. Subsequent PCR with a TSO-specific forward and segment-specific reverse primers was used to differentiate between cRNA with and without a 5ʹ extension. Gel electrophoresis of PCR products resolved faster-migrating bands (cRNA without a 5ʹ extension) from slower-migrating bands (ccRNA with a 5ʹ capped extension). **b**, TSO-based RT-PCR analysis of A549 cells infected with wild-type or T677A mutant viruses at an MOI of 0.3 or mock-infected for 24 hours. Total RNA was either treated or not with oligo d(T)_25_ beads. RT was done with Tuni-13 LNA3 primer (black, for cRNA and ccRNA) or an oligo d(T)_20_ primer (red, for mRNA). **c**, Quantification of fold change in ccRNA and mRNA levels produced by T677A virus as compared to wild-type virus as shown in panel **b**. ccRNA levels were quantified from samples post-oligo d(T)_25_-bead treatment, and mRNA levels were quantified from samples before oligo d(T)_25_-bead treatment. **d**, TSO-based RT-PCR analysis (as in **b**) of RNA extracted one day post-infection from lungs of three mice infected with A/Vietnam/1203/04 (H5N1) and one mock-infected mouse. **e**, Analysis of segment eight (NS) 5ʹ cRNA extensions by RNA circularization, RT-PCR and long-read ONT sequencing for three biological replicates. HEK293-luc cells were infected with wild-type or T677A mutant A/WSN/33 virus at an MOI of 3 for 6 hours. The length of 5ʹ cRNA extension is shown as a distance between cRNA 3ʹ and 5ʹ termini and expressed as proportion of total of cRNA and reads with extensions between cRNA 3ʹ and 5ʹ. **f**, Levels of ccRNAs with 8-30 nt-long capped primer extensions in wild-type and T677A virus infections. **g**, A schematic of ccRNAs with extended internal A-stretch (defined as more than 6 As) and quantification of ccRNA and cRNA reads with an extended A-stretch shown as percentage of all ccRNA with 8-30 nt-long capped primer or of all cRNAs respectively. All gel images are representative of three independent replicates. Data in panel **c, f** and **g** are presented as mean ± SD from three independent experiments. Statistical significance in **f** was determined by an unpaired two-tailed t-test, in **g** by two-way Anova with Tukey’s multiple comparisons test and in **c** by a one-sample t-test where each condition was compared to a theoretical mean of 1 (grouping is for visual purposes only, no comparisons have been made between groups) (ns=non-significant, \**P*<0.05, \*\**P*<0.01, \*\*\**P*<0.001, \*\*\*\**P*<0.0001).

For ccRNA detection during infection we used wild-type and T677A A/WSN/33 (H1N1) virus stocks that were plaque-purified to minimize DelVG content and sequence-verified using Sanger sequencing (Extended Data Fig. 2c). To mitigate unspecific mRNA amplification by the TSO-based RT-PCR, total RNA from wild-type and T677A virus-infected A549 cells was treated with oligo d(T)_25_ beads, which removed approximately 95% of viral mRNAs as confirmed by RT-qPCR (Extended Data Fig. 2d). Subsequent RT-PCR analysis with a terminal cRNA 3ʹ-specific RT primer revealed two bands for all segments: a lower cRNA band and an upper ccRNA band (Fig. 2b). RT-PCR analysis using an oligo d(T)_20_ RT primer showed a single mRNA band. The oligo d(T)_25_ bead treatment did not affect cRNA levels as expected (Extended Data Fig. 2e). The mRNA signal was significantly more reduced after oligo d(T)_25_-bead depletion in comparison to the ccRNA signal for all segments except segment 4 (HA) of the wild-type virus, indicating that mRNA contributed minimally to the ccRNA signal in infection samples after poly(A) RNA depletion (Extended Data Fig. 2f). Consistent with the *in vitro* RNP assays, the T677A mutation increased ccRNA levels for six of the eight segments and the mRNA levels of T677A virus were comparable or slightly reduced relative to the wild-type virus for seven of the eight segments (Fig. 2c). Finally, we confirmed that the T677A mutation did not increase the synthesis of non-canonical replication products, such as mvRNAs and DelVGs (Supplementary Fig. 2a, b).

Using TSO-based RT-PCR we also detected ccRNAs for all eight segments in RNA samples obtained from murine lungs infected with the wild-type A/Vietnam/1203/04 (H5N1) virus (Supplementary Table 1). cRNA levels were unaffected after oligo d(T)_25_-bead treatment as expected and mRNA levels were significantly more reduced in comparison to ccRNA levels indicating that the ccRNA detection was specific (Fig. 2d, Extended Data Fig. 2g, h). No signals were detected in the mock-infected control. These data thus indicate that ccRNA molecules are produced in *in vitro* and *in vivo* infections by wild-type viruses, and that the T677A mutation increases ccRNA production.

To further confirm that the ccRNAs can be produced from a full-length segment in infection, we used RNA circularization and long-read sequencing with Oxford Nanopore Technology (ONT). Total cellular RNA extracted 6 hours post-infection was treated with oligo d(T)_25_ beads to deplete mRNAs and circularized using T4 ssRNA ligase after enzymatic removal of m^7^G caps and pyrophosphates. Segment 8-specific primers were then used to reverse-transcribe and PCR-amplify circularized positive-sense RNAs across the junction between the 5ʹ and 3ʹ termini. After PCR amplification bands were gel-purified and sequenced using ONT (Extended Data Fig. 3a). To control for and quantify template switching and intermolecular ligation events between cRNAs and mRNAs, we designed a segment 8-based spike RNA that contained two barcodes adjacent to the RT and PCR primer binding sites, and two barcodes at the spike termini (resulting in two barcodes at the opposite ends of the resulting amplicon and two at the ligation point, Extended Fig. 3a, Supplementary Fig. 6). This spike RNA was *in vitro* transcribed and added in excess to the ligation reaction. By calculating the proportion of reads with mixed spike and viral primer-adjacent barcodes, we estimated that the template switching frequency between the extreme ends of the RNA molecules was below 25% (Supplementary Table 2). Although the samples contained one to three times the amount of spike RNA to viral RNA, the rate of intermolecular ligation (estimated as the number of reads with a cRNA 3ʹ end ligated to a spike 5ʹ end) was under 0.1% for all samples (Supplementary Table 3). ccRNA sequences were identified based on the schematic shown in Supplementary Fig. 7 (see also Supplementary Note).

The analysis of nucleotide insertions between the 3ʹ and 5ʹ cRNA termini of circularized RNAs showed three peaks for both wild-type and T677A virus infected samples. The first two peaks had a similar maximum for both viruses and consisted of AA or TA dinucleotides (2-nt peak) or a direct repeat of the first nucleotides of the 5ʹ cRNA sequence (6-nt and 7-nt peaks), indicating that they likely originate from realignment events, which have been observed during mRNA synthesis^20,36^ and here likely occurred during replication initiation. The third peak consisted of 8 to 30 nt-long random sequence insertions and had a maximum at 11-12 nt (Fig. 2e, Extended Data Fig. 3b), which is consistent with the mean length of capped host primers produced by the IAV polymerase and thus represents true ccRNAs in our dataset^20^. This third peak showed that of all reads containing 3ʹ and 5ʹ cRNA termini (i.e., all cRNAs and ccRNAs) the ccRNA content was ∼1% for the wild-type virus, while this number increased to 18-20% for the T677A mutant (Fig. 2f). The proportion of ccRNA reads for both wild-type and T677A mutant viruses was significantly higher than the rate of intermolecular ligation, suggesting that ligation between cRNA and mRNA molecules, as opposed to ccRNA circularization, was negligible (Extended Data Fig. 3c, Supplementary Table 3). Our sequencing analysis also revealed that 5-7% of wild-type virus ccRNAs and 10-12% of T677A virus ccRNAs had an extended internal A-stretch upstream of the cRNA 3ʹ terminus that was longer than the vRNA 5ʹ U-stretch, suggesting that internal polyadenylation was attempted by the RNA polymerase during transcription termination. In contrast, only around 1% of cRNAs had an extended internal A-stretch (Fig. 2g, Supplementary Table 4). This finding shows that some ccRNAs are generated when the polymerase fails to complete polyadenylation and instead returns to copying the cRNA 3ʹ terminus and that the coupling between replication/transcription initiation and termination is error prone even for the wild-type RNA polymerase. We confirmed that the samples used for ONT sequencing showed increased ccRNA production by the T677A mutant in comparison to the wild-type virus using the TSO-based RT-PCR (Extended Data Fig. 3d).

Our analysis also showed differences in the 5ʹ terminus of cRNA molecules, with around 5% of T677A virus cRNAs lacking the first adenosine, in comparison to around 1% of wild-type virus cRNAs (Extended Data Fig. 3e). Initiation of cRNA synthesis from the second instead of the first nucleotide of the template has been documented before *in vitro*^37,38^ and here we show evidence of this initiation mode in infection and that the PB1 T677A mutation affects this allosterically. Overall, these data show that the ccRNA synthesis occurs in infections by different IAV strains in both cell culture and in animal infection models, and that perturbation of transcription termination increases ccRNA synthesis.

### T677A mutation and ccRNA synthesis correlate with enhanced IFN responses

As the T677A mutant virus was previously shown to stimulate IFN responses in infected cells^15^ we measured IFN induction by wild-type and T677A mutant viruses in an infection time course. The T677A virus caused significantly higher *IFN-β* promoter activation than the wild-type virus at 12 and 24 hours post-infection in IFN-β luciferase reporter HEK293 cells (HEK293-luc) (Fig. 3a). To determine if the *IFN-β* promoter activation was RIG-I-dependent, we used *RIG-I* -/- and *MAVS* -/- HEK293-luc cells^39^ in which knock-out efficiency was validated by western blot and IFN-β luciferase reporter functionality was verified by MAVS overexpression (Extended Data Fig. 4a, b). Additionally, by transfecting HMW poly(I:C), we verified that MDA-5 signalling remained intact in *RIG-I* -/- cells but not *MAVS* -/- cells, while both knock-out cells lost the ability to respond to RIG-I ligands such as LMW poly(I:C) and *in vitro* transcribed, gel-purified NA196 vRNA. In contrast, all cell lines retained functional DNA sensing pathways as they responded to HSV-1 60-mer DNA transfection (Extended Data Fig. 4c). Using these wild-type and knock-out cells we showed that T677A virus induced higher *IFN-β* promoter activity and interferon stimulated genes (ISG) expression than the wild-type virus in wild-type HEK293-luc cells, while both viruses failed to activate IFN or induce ISG expression in *RIG-I* -/- or *MAVS* -/- cells (Fig. 3b, c). Similar RIG-I-dependent ISG induction was observed for both wild-type and T677A mutant viruses in A549 cells (Extended Data Fig. 4d), confirming that the T677A virus activates IFN responses via the RIG-I pathway. The T677A virus also exhibited reduced growth in HEK293-luc and A549 cells and formed smaller plaques in MDCK cells compared to the wild-type virus. The growth impairment was evident in both wild-type and immunodeficient HEK293-luc and A549 cells, indicating that the T677A mutation affects virus growth regardless of its impact on immune activation (Fig. 3d; Extended Data Fig. 4e, f).

**Fig. 3.**
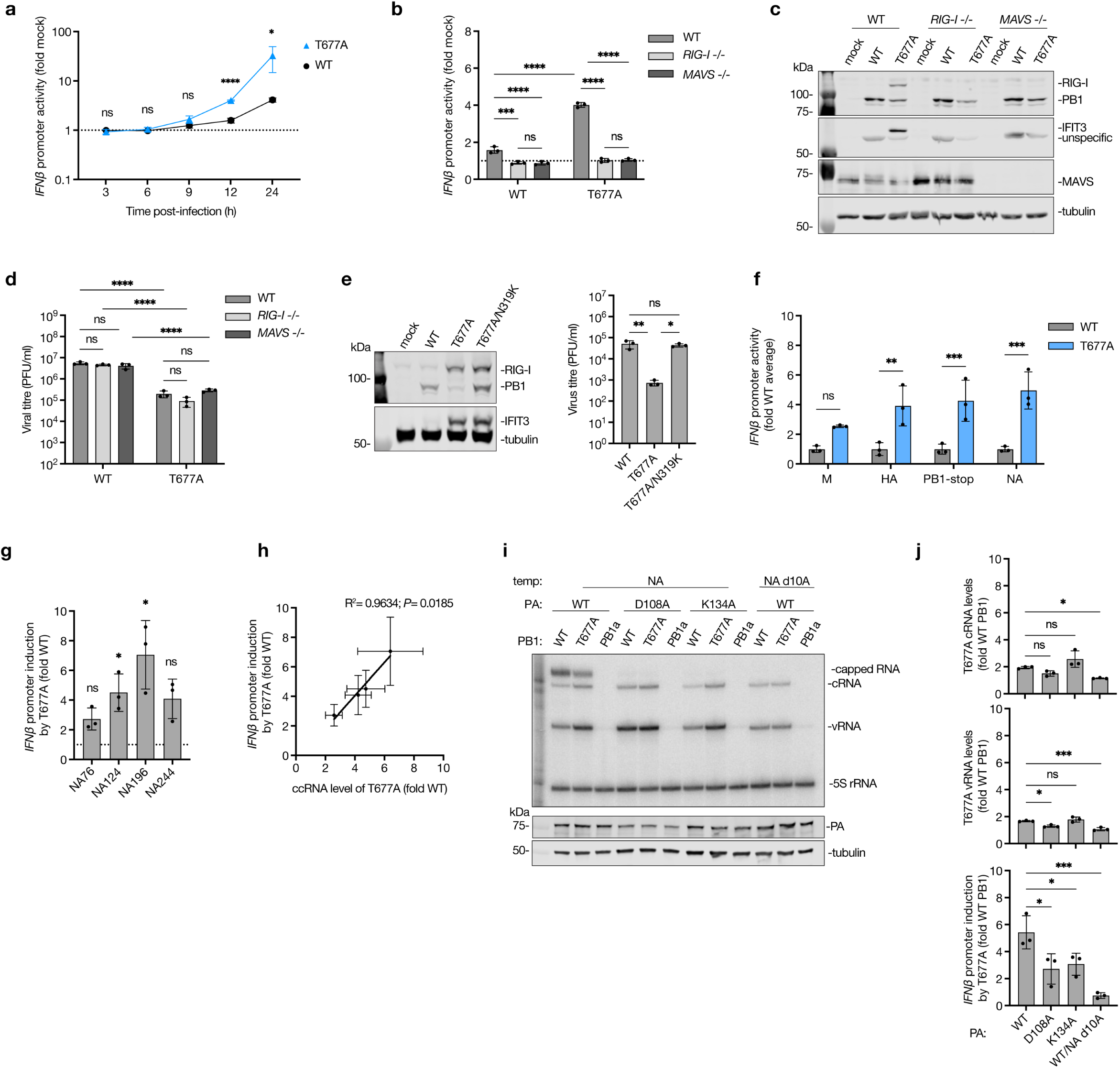
The effect of T677A mutation on innate immune activation in infection and RNP assay. **a**, HEK293-luc cells were infected with either wild-type or T677A mutant A/WSN/33 viruses at an MOI of 3, or mock-infected, in a synchronized infection. IFN-β luciferase reporter activity was measured at various time points post-infection. **b**, Wild-type, *RIG-I* -/- or *MAVS* -/- HEK293-luc cells were infected with wild-type or T677A mutant viruses at an MOI of 3, or mock-infected. IFN-β luciferase reporter activity was measured at 12 hours post-infection. **c**, Wild-type, *RIG-I* -/- or *MAVS* -/- HEK293-luc cells were infected with wild-type or T677A mutant viruses at an MOI of 0.01, or mock-infected for 48 hours. Protein expression was analyzed by western blotting. **d**, Viral titers in cell culture supernatants collected 48 hours post-infection with an MOI of 0.01 as quantified by plaque assay. **e**, HEK293-luc cells were infected with wild-type, T677A mutant or T677A/N319K double mutant viruses at an MOI of 0.01, or mock-infected for 48 hours. Protein expression was analyzed by western blotting, and viral replication was measured by plaque assay. **f**, RNP reconstitution assay with wild-type and T677A PB1, four different full-length segments and IFN-β luciferase reporter was done in HEK293T cells, and *IFN-β* promoter activation was measured 24 hours post-transfection. **g**, *IFN-β* promoter activation by T677A versus wild-type polymerases in RNP reconstitution assay with short segment 6-based templates as described in Fig. 1e. **h**, Correlation between *IFN-β* promoter activation and ccRNA synthesis levels from short segment 6-based templates in the RNP reconstitution assay described in Fig. 1e. **i**, RNP reconstitution assay with wild-type or T677A PB1, wild-type or transcriptionally inactive mutants of PA (D108A and K134A) and segment 6 (NA) template, or wild-type PA and the NA template lacking the 10^th^ adenosine from the 5ʹ terminus (NA d10A). Primer extension was done with internal +sense (NA 160) and -sense (NA 1280) primers, PA expression was analyzed by western blotting. **j**, Quantification of cRNA and vRNA synthesis (capped RNA not quantified due to lack of signal) and *IFN-β* promoter activation by T677A versus wild-type polymerases in the RNP reconstitution assay described in panel i. All gel images are representative of three independent replicates. Data in all graphs are presented as mean ± SD from three independent experiments. Statistical significance was determined using a one-way ANOVA with Dunnett’s multiple comparisons test (**a**, **j**), two-way ANOVA with Šídák’s multiple comparisons test (**b**, **f**) or Tukey’s multiple comparisons test (**d**), one-sample t-test (**g**), and simple linear regression analysis (**h**); (ns=non-significant, \**P*<0.05, \*\**P*<0.01, \*\*\**P*<0.001, \*\*\*\**P*<0.0001).

To explore if the T677A mutant virus could compensate for or recover from the growth defect, we passaged wild-type and mutant viruses in A549 cells and performed whole viral genome sequencing at the passages where phenotypic changes occurred. Two of the three T677A passaging replicates (2 and 3) reached wild-type virus titres and wild-type ISG expression levels at early passages by reverting to the wild-type threonine at position 677. Replicate one did not revert to wild-type and showed a gradual increase in titre (Extended Data Fig. 4g, Supplementary Tables 5 and 6). Interestingly, in all three replicates we observed a selection for the NP N319K mutation that has previously been shown to enhance adaptation of the avian IAV to mice^40^ (Supplementary Table 6). We characterized the double N319K/T677A mutant in an RNP assay and showed that NP N319K did not affect cRNA nor vRNA synthesis by the T677A mutant, and that the double T677A/N319K mutant generated equally elevated ccRNA levels as the single T677A mutant (Supplementary Fig. 3). However, in infection, the T677A/N319K double mutation restored viral titres to wild-type virus levels even though it induced similar levels of ISG expression as the T677A mutant. This confirmed that the growth impairment of the T677A mutant is independent of its immunostimulatory activity and that it can be rescued by the addition of the NP N319K mutation (Fig. 3e).

Next, we used RNP reconstitution coupled with IFN-β reporter assays to determine if the impaired transcription termination activity of the T677A mutation was sufficient to recapitulate the increase in innate immune activation observed in infection. We found that the viral polymerase containing the T677A mutation markedly increased *IFN-β* promoter activity in comparison to the wild-type polymerase in the RNP assays with four different full-length templates and the short segment 6-based templates (Fig. 3f, g). We also observed a positive correlation between the *IFN-β* promoter activity and the ccRNA levels generated on the segment 6-based templates (Fig. 3h). However, neither of these outcomes was strictly associated with the template length (Fig. 1e, 3g). At the same time, both ccRNA levels and IFN induction were low on all segment 5-based templates of different lengths (Supplementary Fig. 4a, b), suggesting that ccRNA production is likely influenced by the template sequence and/or secondary structure, rather than length.

To test whether immunostimulatory activity of the T677A mutation is dependent on viral transcription, we used an RNP assay with cap-snatching-deficient PA mutants (K134A and D108A)^41^ or a segment 6 template lacking the 10th adenosine (NA d10A) from the 5ʹ vRNA terminus, which abolishes transcription initiation but not replication^42^ (Fig. 3i). While some conditions also affected replication, all conditions abrogated capped RNA synthesis and significantly diminished IFN activation by the T677A mutant (Fig. 3i, j). We also showed that T677A mutation did not destabilise polymerase-vRNA binding, which could have made it available for RIG-I recognition (Supplementary Fig. 5). Together, we show that ccRNA synthesis contributes to innate immune activation in an RNP assay and in infection.

### RIG-I binds ccRNA in an RNP assay and infection

To investigate whether ccRNA molecules are recognised by RIG-I, we co-expressed Myc-tagged RIG-I or its 5ʹ-triphosphate-binding mutant (K861A/K858A/K851A) in an RNP assay with a full-length segment 6 template in HEK293T cells. Subsequent RIG-I immunoprecipitation and primer extension with internal primers showed increased vRNA and cRNA levels in both total and immunoprecipitated fractions in the presence of the T677A mutant polymerase in comparison to the wild-type polymerase. The T677A mutation also increased NP but not PB1 co-immunoprecipitation with RIG-I, indicating polymerase-depleted RNP binding. Furthermore, there was a strong enrichment of capped RNA binding to RIG-I in the presence of the T677A polymerase (Fig. 4a). TSO-based RT-PCR analysis confirmed that the capped RNA molecules observed in the primer extension reactions represented ccRNAs (Fig. 4b). In contrast, no significant enrichment in viral RNA or NP was observed with T677A polymerase on the full-length segment 5 template (Extended Data Fig. 5a), in line with the lower levels of ccRNA synthesis and IFN upregulation by the T677A polymerase on the short segment 5-based templates (Supplementary Fig. 2a, b).

**Fig. 4.**
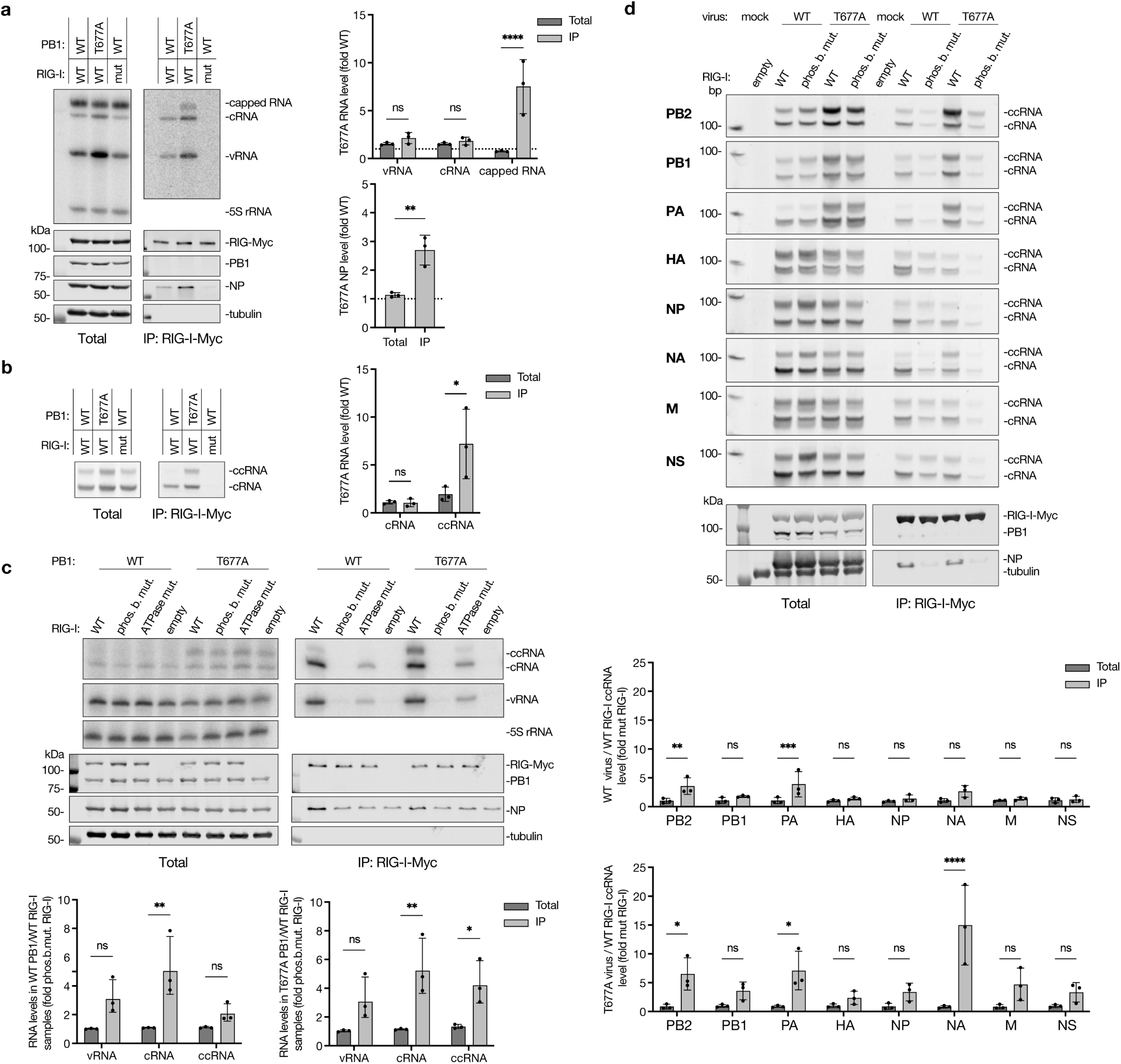
ccRNA binding by RIG-I. **a**, Myc-tagged RIG-I was co-expressed with wild-type or T677A mutant RNP and segment 6 template in HEK293T cells, followed by immunoprecipitation. A phosphate-binding mutant of RIG-I was used as a negative control. RNA levels in total and immunoprecipitated (IP) fractions were measured by primer extension with internal +sense (NA 160) and -sense (NA 1280) primers, and protein levels were assessed by western blotting. Shown are a representative image from three independent experiments (left) and quantification (right). **b**, TSO-based RT PCR for segment 6 cRNA and ccRNA was done in total and IP fractions generated as in (a) with quantification. **c**, Immunoprecipitation of wild-type, phosphate-binding mutant, or ATPase mutant Myc-tagged RIG-I was performed in an RNP assay with a short NA196 template. An empty plasmid served as a negative control. RNA levels were assessed by primer extension assay with terminal cRNA (NA 5′) and vRNA (NA-2) primers (quantification is shown) and protein levels assessed by western blotting. The quantification compares the RNA signal in wild-type RIG-I samples to the phosphate-binding mutant RIG-I samples for total and IP fractions. HEK293T cells were transfected with Myc-tagged RIG-I and 24 h later infected with an MOI of 0.1 of wild-type and T677A viruses (wild-type virus inoculum was adjusted to an MOI of 0.1 of T677A virus by qPCR targeting PB1 vRNA to account for variations in incoming RNA levels). RIG-I immunoprecipitation was done 24 h after infection. TSO-based RT PCR was used to determine cRNA and ccRNA levels for each segment in total and IP fractions. Protein levels were assessed by western blot. All gel images are representative of three independent replicates. Data are presented as mean ± SD from three independent experiments. Statistical significance was determined using two-way ANOVA with Šídák’s multiple comparisons test (**a**, top right, **b**, **c**, **d**) or an unpaired two-tailed t-test (**a**, bottom right) (ns=non-significant, \**P*<0.05, \*\**P*<0.01, \*\*\**P*<0.001, \*\*\*\**P*<0.0001).

RIG-I immunoprecipitation with a short NA196 template also showed cRNA, vRNA and ccRNA enrichment in RIG-I-bound fractions for both wild-type and T677A mutant polymerases, and the T677A mutation resulted in a stronger ccRNA enrichment than the wild-type polymerase (Fig. 4c, Extended Data Fig. 5b). Nucleoprotein, but not PB1, also co-immunoprecipitated with the wild-type RIG-I in both wild-type and T677A polymerase conditions, indicating polymerase-depleted RNP binding. Both phosphate-binding and ATPase (K270A) mutations in RIG-I reduced precipitation of all viral RNA species, underscoring the necessity of these activities for RIG-I-mediated RNA detection, including ccRNA binding (Fig. 4c, Extended Data Fig. 5b).

Next, we tested whether RIG-I binds ccRNA molecules produced in infection. RIG-I immunoprecipitation 24 hours post-infection with an MOI of 0.1 of wild-type and T677A mutant viruses showed segment-specific ccRNA enrichment in wild-type RIG-I-bound fractions in comparison to phosphate-binding mutant RIG-I. There was a significant enrichment of PB2 and PA ccRNAs for both viruses and NA ccRNA for the T677A mutant virus (Fig. 4d). NP and cRNAs were also enriched in immunoprecipitated samples for both viruses as expected (Fig. 4d, Extended Data Fig. 5c). These data showed that ccRNAs produced by both wild-type and to a larger extent T677A mutant viruses are detected by RIG-I during infection.

### RIG-I is activated by ccRNA in complex with a negative-sense viral RNA

Given that the cap-1 structures present on IAV cap-snatched primers are incompatible with RIG-I activation^43^, we assessed whether ccRNA molecules alone are immunostimulatory. To this end, we used NA196-based RNAs that were produced by *in vitro* transcription and gel-extraction (Extended Data Fig. 2a) and modified at their 5ʹ termini to mimic different viral RNA species. Capped RNAs – including ccRNA, ccRNA without a 3ʹ terminus, and mRNAs – were capped and any remaining 5ʹ-triphosphorylated RNAs were enzymatically digested to avoid unspecific RIG-I activation. cRNAs and vRNAs were left untreated as they already contained 5ʹ-triphosphates following T7 transcription, like the true viral RNAs. Transfection of *in vitro* transcribed RNAs into HEK293 IFN-β reporter cells showed that cRNA and vRNA molecules induced a strong, dose-dependent *IFN-β* promoter activation as expected, while ccRNA, ccRNA-3ʹ, and mRNA molecules did not, showing that ccRNAs alone do not activate RIG-I (Fig. 5a).

**Fig. 5.**
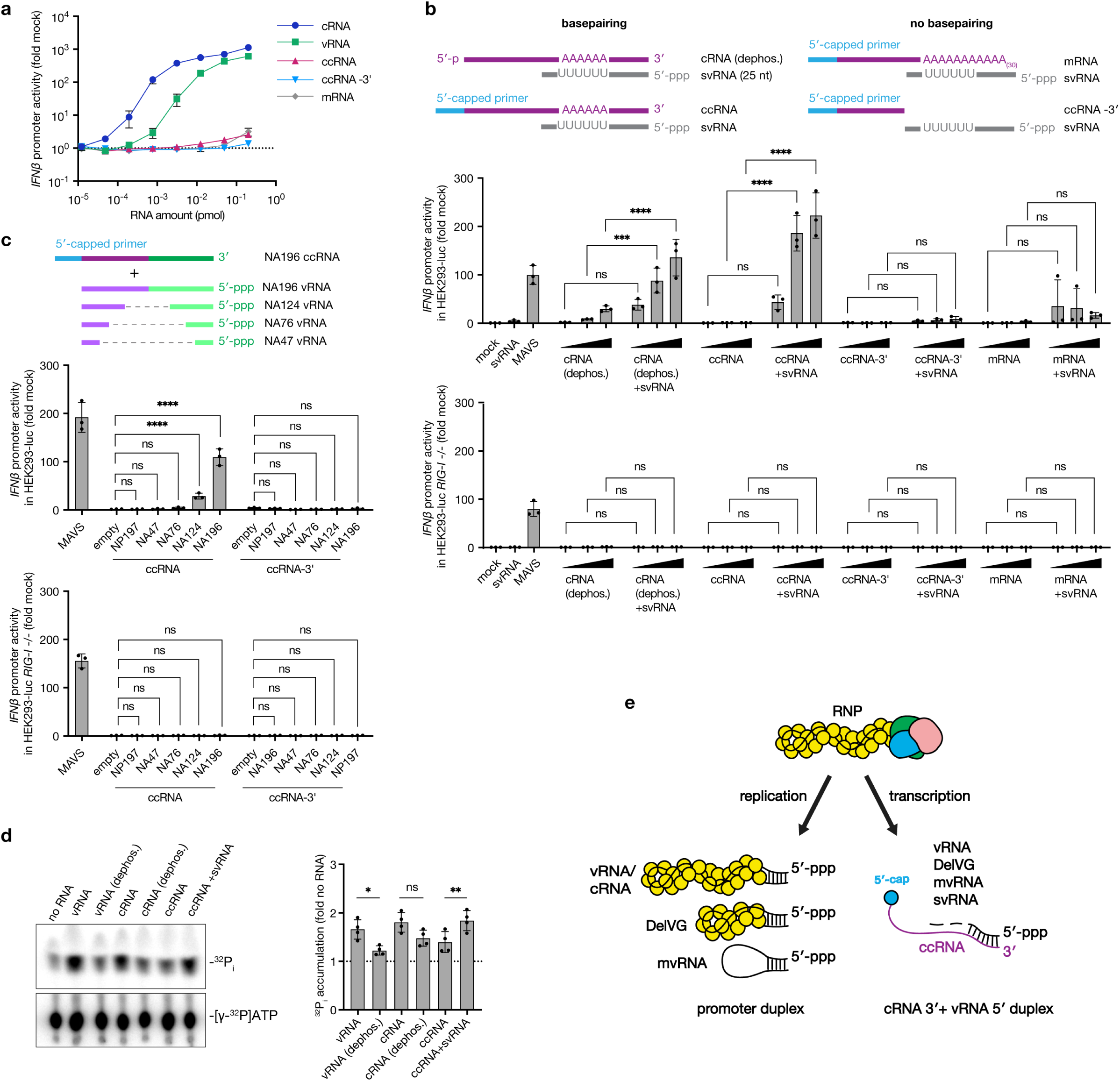
RIG-I activation by ccRNA complexes with complementary negative-sense RNA. **a**, HEK293-luc cells were transfected with a two-fold dilution series of *in vitro* transcribed NA196-based vRNA, cRNA, ccRNA, ccRNA-3ʹ, or mRNA, starting at 0.2 pmol. *IFN-β* promoter activity was measured 20 hours post-transfection. **b**, One pmol of segment 6 svRNA was transfected alone or with increasing concentrations (0.05, 0.1 and 0.2 pmol) of *in vitro* transcribed NA196-based dephosphorylated cRNA, ccRNA, ccRNA-3ʹ and mRNA into HEK293-luc wild-type or *RIG-I* -/- cells. Total HEK293T RNA (mock) was used as a negative control and MAVS-encoding plasmid was used as a positive control. *IFN-β* promoter activity was measured 20 hours post-transfection. Schematic shows expected complementary regions between svRNA and the 3ʹ terminus of cRNA/ccRNA or the conditions where terminal base pairing doesn’t occur. **c**, 0.2 pmol of *in vitro* transcribed NA196-based ccRNA or ccRNA-3ʹ or equivalent amount of total HEK293T (mock) RNA was co-transfected with pPolI plasmids encoding 47-196 nt long segment 6-based vRNA templates or segment 5-based NP197 template in HEK293-luc wild-type or *RIG-I* -/- cells. An empty pPolI plasmid was used as a negative control, while MAVS-encoding plasmid was used as a positive control. *IFN-β* promoter activity was measured 20 hours post-transfection. The schematic shows the NA196 ccRNA with purple and green colours indicating segment 6 sequences on either side of the internal deletion. Complementary vRNA sequences are shown in lighter colours and internal deletions relative to NA196 template are indicated with a dashed line. **d**, ATPase activity of recombinant RIG-I was assessed in the presence of NA196 ccRNA or its complex with segment 6 svRNA. NA196 vRNA and cRNA were used as positive controls and their dephosphorylated versions as negative controls. ^32^P_i_ release/accumulation is used as a measurement of ATPase activity. A representative image from four independent experiments is shown. **d**, A proposed model of RIG-I activation during infection. During viral replication RIG-I can detect partially complementary triphosphorylated promoters of vRNAs. cRNAs, DelVGs or mvRNAs in the polymerase-unbound form. During transcription the 3ʹ terminus of ccRNA, can base pair with the complementary 5ʹ-triphosphorylated terminus of a negative-sense RNA resulting in a double-stranded 5ʹ-triphosphorylated ligand for RIG-I. Data in panels **a**-**d** are presented as mean ± SD from three (a-c) or four (d) independent experiments. Statistical significance was determined using two-way ANOVA with Šídák’s multiple comparisons test (**b**, **c, d**) (ns=non-significant, \**P*<0.05, \*\**P*<0.01, \*\*\**P*<0.001, \*\*\*\**P*<0.0001).

We hypothesised that during infection ccRNAs might be detected by RIG-I when their 3ʹ terminus is base-paired with their complementary negative-sense 5ʹ-triphosphorylated RNA, which could be a vRNA, DelVG, mvRNA, or svRNA of the same segment. To test this hypothesis, we co-transfected positive-sense NA196-based RNAs with a negative-sense svRNA, which consisted of twenty-five 5ʹ-terminal nucleotides of segment 6 vRNA and a 5ʹ-triphosphate. MAVS-FLAG transfection was used as a positive control and it induced IFN as expected, while svRNA alone did not. When molecules containing the cRNA 3ʹ terminus, such as cRNA (dephosphorylated, to avoid cRNA promoter-mediated activation) or ccRNA were co-transfected with svRNA it resulted in dose-dependent immune activation. In contrast, co-transfection of svRNA with molecules that lacked the cRNA 3ʹ terminus, such as mRNA and ccRNA-3ʹ did not induce IFN, indicating that the presence of the cRNA 3ʹ terminus is driving ccRNA-mediated immune activation as it base pairs with its complementary svRNA. No IFN induction was observed when RNAs were co-transfected in *RIG-I* -/- HEK293-luc cells, demonstrating that their detection is mediated by RIG-I (Fig. 5b).

We also tested if NA196 ccRNAs could activate RIG-I in the presence of complementary vRNA templates of the same length or shorter. The vRNA templates had different sizes of internal deletions but conserved 5ʹ and 3ʹ termini. Two longer vRNA templates, but not the two shorter ones, induced *IFN-β* promoter activation when added to ccRNA. Co-transfection of NA196 ccRNA with segment 5 based NP197 template did not activate IFN indicating that template complementarity is required. Similarly, NA196 ccRNA-3ʹ did not induce *IFN-β* promoter activation with any of the vRNA templates showing the importance of 3ʹ-terminal nucleotides. The ccRNA-vRNA complexes failed to induce IFN in *RIG-I* -/- HEK293-luc cells, confirming that their detection was RIG-I-dependent (Fig. 5c).

Finally, we tested the ability of ccRNA molecules to activate purified RIG-I *in vitro* by measuring ATPase activity. vRNA and to a lesser extend cRNA transcripts showed RIG-I activation in comparison to their de-phosphorylated versions as expected. In support of our previous results, RIG-I activation by ccRNA was increased significantly in the presence of svRNA (Fig. 5d). Overall, these findings indicate that ccRNAs can contribute to collective RIG-I activation provided that they hybridise with a complementary negative-sense 5ʹ- triphosphorylated RNA.

## Discussion

It is well-established that IAV genomes and non-canonical replication products are detected by RIG-I during viral infection (Fig. 5e). However, several studies in which replication was blocked or transcription enhanced suggested that IAV transcription also contributes to innate immune activation during infection^29–33^. In this study, we identified non-canonical transcription products, ccRNAs, which are produced during aberrant transcription termination and contribute to the innate immune activation (Fig. 5e). ccRNAs are capped transcripts that contain a cRNA 3ʹ terminus instead of a poly(A) tail. We show that ccRNAs are produced by the viral polymerase *in vitro*, in RNP assays, in cell culture, and in murine infection models. We further show that ccRNAs are detected by RIG-I in both RNP assays and during infection with wild-type and T677A mutant viruses. We propose a mechanism by which ccRNAs, although not inherently immunostimulatory, can form RIG-I ligands when base paired with a negative-sense RNA. Since ccRNAs are produced during transcription, just like mRNAs, they are unlikely to associate with viral NP or polymerase and therefore are available to hybridise with other unencapsidated negative-sense RNAs such as svRNAs or short mvRNAs (Fig. 5e). It is also possible that a ccRNA could base pair with its template vRNA during transcription termination to stimulate the release of the template, as we recently proposed for mvRNAs^44^. This mechanism might also explain innate immune activation during primary transcription^29–32^, when the negative-sense template is only present in the form of an RNP. Even though cRNAs possess the same 3ʹ cRNA terminus as ccRNAs, they are unlikely to activate RIG-I by the same mechanism since they are encapsidated by the viral polymerase and NP. In our study, ccRNAs contributed significantly to the immunostimulatory activity of the T677A mutant virus yet their relative contribution to RIG-I activation compared to other immunostimulatory RNA species in wild-type infection and disease remains to be determined.

The effect of the T677A mutation on polyadenylation provides a new insight into transcription termination and into the function of the triple-stranded β-sheet - a key structural feature of the active IAV RNA polymerase and one that is less prominent in other negative-sense RNA virus RNA polymerases^23,45^. It is currently unclear what factors determine whether the polymerase commits to stuttering on the U-stretch during transcription or to copying the vRNA 5ʹ terminus during cRNA synthesis. Our sequencing results show an increase in A-stretch extensions in ccRNAs compared to cRNAs, suggesting that the viral polymerase can switch between polyadenylation and replication during transcription termination. ccRNAs represented only 1% of all sequences containing the cRNA 3ʹ and 5ʹ termini during wild-type virus infection, however the T677A mutation in the triple-stranded β-sheet increased ccRNA generation nearly 20-fold as well as the occurrence of A-stretch extensions. While this manuscript was in revision, a mutagenesis study showed that other mutations in the triple-stranded β-sheet region (of PB1 residues L675 and N676) impair polyadenylation *in vitro*, producing readthrough products that are likely similar in nature to the ccRNAs characterized in this study^46^. Overall, these lines of evidence indicate that the formation of the triple-stranded β-sheet is critical for canonical mRNA synthesis.

As with other non-canonical IAV RNAs, the potential evolutionary advantage of ccRNA production for the virus remains unclear. Although the growth of the T677A mutant virus was impaired in cell culture, it could be restored to wild-type virus levels by the addition of the N319K mutation in NP, which did not affect ccRNA levels or innate immune activation, indicating that ccRNAs per se are not inhibitory in cell culture conditions. Thus, their potential function, cellular localization, translational potential, and stability in the cell remain an open question. Overall, with the recent discoveries of svRNAs, mvRNAs, and now ccRNAs, our findings suggest that the repertoire of non-canonical IAV RNAs may be far more extensive and complex in their interactions than previously assumed. Further exploration of the non-canonical RNA diversity, along with their roles in viral life cycle and pathogenesis, may be essential for our understanding of influenza disease and the development of more effective antiviral strategies.

## Methods

### Cells

Human embryonic kidney cells (HEK293T) and Madin-Darby canine kidney cells (MDCK) were sourced from the American Type Culture Collection. Wild-type, *RIG-I* -/- and *MAVS* -/- human embryonic kidney cells stably expressing firefly luciferase and GFP under the *IFN-β* promoter (HEK293-luc) were a generous gift from J. Rehwinkel (Oxford University)^39^. Wild-type and *RIG-I* -/- human alveolar basal epithelial cells (A549) were kindly provided by B. Ferguson (Cambridge University)^47^. Cells were cultured in Dulbecco’s Modified Eagle’s Medium (DMEM, Sigma-Aldrich) supplemented with 10% foetal calf serum (FCS, Gibco) and 2 mM L-glutamine (Gibco). We will refer to this medium as complete DMEM.

### Plasmids

PB1 mutations described in this study, NP N319K mutation, RIG-I K270A and K861A/K858A/K851A mutations were introduced using site-directed mutagenesis using primers listed in Supplementary Table 7. The pPolI plasmids encoding A/WSN/33 genome segments were previously described in^48^; the pPolI plasmids encoding segment 5- and segment 6-based templates with internal deletions were described in^26^; the pPolI-NA d10A template was described in^42^; the pcDNA3 plasmids encoding wild-type A/WSN/33 RNP proteins in^49^; pcDNA3 plasmids encoding A/Brevig Mission/1/1918 H1N1 (BM18), A/Fuijan/1/1995 H5N1 and A/Vietnam/1203/2004 H5N1 polymerases in^26^; pcDNA3 plasmids encoding A/Bilthoven/16190/1968 H3N2 and A/Netherlands/499/2017 H3N2 polymerases in^50^; the pcDNA3 plasmid encoding PB1 D445A/D446A (PB1a) mutant in^51^; the pcDNA3 plasmid encoding PB1 R238A/R672A mutant in^52^; the pcDNA3 plasmid encoding PA with the D108A or K134A mutations in^41^; the pcDNA3 plasmid encoding TAP-tagged PB2 in^53^; the plasmid expressing firefly luciferase under *IFN-β* promoter (pIFD(−116)lucter) and pcDNA3 encoding *Renilla* luciferase in^26^; the pcDNA plasmid encoding wild-type Myc-RIG-I in^26^; MAVS-FLAG encoding plasmid and corresponding empty plasmid in^54^.

### Animal infections

Recombinant A/Vietnam/1203/04 (H5N1) virus was generated as described previously^55^. Recombinant virus was passaged in MDCK cells at an MOI of 0.001 before use in *in vivo* experiments. Work with highly pathogenic H5N1 virus was conducted in the Biosafety Level 3 laboratory at the LKS Faculty of Medicine, The University of Hong Kong following approved guidelines and ethics approved by the Committee on the Use of Live Animals in Teaching and Research (CULATR). BALB/c mice were infected intranasally with 10^5^ PFU of A/Vietnam/1203/04 (H5N1) virus or PBS control and sacrificed 1 day post infection. Weights were recorded and virus titres in the lungs were determined using plaque assay on MDCK cells.

### Generation of recombinant viruses

Wild-type, T677A and T677A/N319K mutant A/WSN/33 viruses were generated by reverse genetics using the 12-plasmid system. Briefly, eight pPolI plasmids encoding the eight IAV genome segments and four pcDNA3 plasmids encoding the IAV PB2, PB1, PA and NP proteins were transfected into HEK293T cells using Lipofectamine 2000 (Invitrogen) in 6 cm dishes and grown in complete DMEM. Twenty-four hours later, the cell culture medium was replaced by DMEM supplemented with 0.5% FCS and 2 mM L-glutamine. We will refer to this medium as the infection DMEM. After 48 h, cell-culture supernatants were collected, cleared by centrifugation at 1,000 rpm for 5 min, and used to infect a monolayer of MDCK cells in T75 flasks. Forty-eight hours later, cell culture supernatants were collected and cleared by centrifugation at 2,000 rpm for 10 min. The initial virus stocks were plaque-purified on MDCK cells, and viruses grown from individual plaques were subsequently assessed for DelVG content using RT-PCR. Virus isolates containing the lowest amount of DelVGs were selected and used for downstream experiments. Mutations in PB1 and NP segments were validated with Sanger sequencing, and viral titres were determined by plaque assay using MDCK cells.

### Virus infection

For infection experiments, cells were seeded in 24-well plates at 2*10^5^ cells/well in complete DMEM. For infection of HEK293-luc cells, wells were pre-treated with 0.5 mg/ml poly-L-lysine (Sigma-Aldrich). The next day, cells were infected with an MOI indicated for specific experiments in infection DMEM at 37 °C, 5% CO_2_ for 1 h. After infection, cells were washed with PBS and incubated in infection DMEM at 37 °C, 5% CO_2_. At the time periods indicated for specific experiments, supernatants were collected for virus titre quantification by plaque assay, and cells were lysed in either TRI Reagent (Sigma-Aldrich) for phenol-chloroform RNA extraction and isopropanol precipitation as described previously^56^ or Laemmli buffer for western blotting.

For synchronised infections, HEK293-luc cells were seeded at 4*10^4^ cells/well in a 96-well plate pre-treated with poly-L-lysine. The next day, cells were infected with an MOI of 3 at 4 °C for 1 h. Cells were then washed with PBS and incubated in infection DMEM at 37 °C, 5% CO_2_. At 3, 6, 9, 12 and 24 h time points the cell culture medium was aspirated and the firefly luciferase signal measured using DualGlo reagent (Promega) and a GloMax microplate reader (Promega) or Synergy LX plate reader (BioTek).

### Serial passaging of viruses

To passage viruses A549 cells were seeded at 4*10^5^ cells per well in 12-well plates. The next day cells were infected in triplicate with wild-type or T677A mutant A/WSN/33 viruses at an MOI of 0.01. Separate plates were used for each virus to avoid cross-contamination. Cell culture medium was collected after 48 h and cell debris were removed by pelleting at 13,000 g for 5 min. Supernatants were collected for virus quantification by plaque assay and RNA isolation using TRI Reagent (Sigma-Aldrich). Cell monolayers were lysed with Laemmli buffer for western blotting. Viral titres were measured after each passage, and the titre of each new passage was used for the subsequent infection to create the next passage. A total of seven passages were performed.

### Sample preparation for NGS of passaged viruses

Viral RNA collected from cell culture supernatants was isolated from Tri Reagent using phenol-chloroform extraction and isopropanol precipitation as described previously^56^. The RNA quality was determined using a Qubit RNA high sensitivity kit (Invitrogen). Eight ng of RNA was used for RT with SuperScript III (Invitrogen) and the Tuni-12 primer. PCR amplification was done using *Pfu* DNA polymerase (Promega) and the Tuni-13 and Tuni-12 primers for 30 cycles (Supplementary Table 7). The PCR products were purified using the Monarch PCR and DNA Cleanup Kit (NEB), and the DNA concentration and purity were assessed using the Qubit dsDNA broad range kit (Invitrogen). Amplicons were pooled for each sample and next cleaned up with 0.56x Ampure XP beads to remove short DNA fragments. Next, the amplicons were sheared using a Covaris LE220 sonicator to around 300 bp. Illumina sequencing adapters and barcodes were added to the sheared DNA fragments using a NEBNext Ultra II DNA Library Prep Kit for Illumina (NEB, MA). The purified DNA library products were examined on Bioanalyzer DNA High Sensitivity chips (Agilent, CA) for size distribution, quantified by Qubit fluorometer (Invitrogen, CA), and pooled at equal molar amount. The library pool was sequenced on Illumina MiSeq using a Nano 300 cycle kit as pair-end 150 nt reads. Raw sequencing reads were filtered by the MiSeq control software and only the Pass-Filter (PF) reads were used for further analysis.

### Data processing pipeline of NGS for viral passaging

NGS data were analysed using the Galaxy Princeton Genomics server (galaxy.princeton.edu)^57^. First, TrimGalore!^58^ was used to remove adaptor contamination, trim low-quality ends from reads with a phred quality score below 30, and filter out reads shorter than 100 nt. Read quality assessment was performed using FastQC^59^. The reference genome was based on the plasmid sequences that were used to produce viral stocks. All plasmids’ sequences were confirmed using Sanger sequencing prior to read alignment. The reads were aligned to the reference genome using BWA-MEM^60^. Duplicate sequences were removed using MarkDuplicates with the following parameters: scoring strategy for non-duplicate reads sum of base qualities, validation strategy - lenient. Next, the BAM dataset was filtered using Filter^61^ and only those reads were retained that matched the following parameters: mapping quality ≥ 20, ‘paired’, and ‘properly paired’. BamLeftAlign was used to left-align reads to homogenise the positional distribution of indels. Variant detection was done using FreeBayes^62^ with the following parameters: minimum coverage to process a site: 100; set ploidy for analysis to 1. Only variants with a frequency over 5% were scored.

### Western blotting

Cells were lysed in Laemmli buffer (62.5 mM Tris-HCl pH 6.8, 10% glycerol, 2% SDS, 100 mM DTT (Invitrogen), 0.01% Bromophenol Blue) and sonicated. Proteins were separated using 8% SDS-PAGE and immobilised on a 0.45 µm nitrocellulose membrane (GE healthcare). The membranes were incubated in blocking buffer (PBS, 5% bovine serum albumin (RPI), 0.1% Tween-20 (RPI)) for 1 h, washed 3x with PBST (PBS, 0.05% Tween-20), and then incubated with the primary antibody diluted in blocking buffer for approximately 16 h at 4 °C. The following primary antibodies were used: PB1 (GTX125923, GeneTex), NP (GTX125989, Genetex), PA (GTX125932, Genetex), tubulin (MCA776, Serotec), IFIT3 B7 (sc-393512, Santa Cruz), IFIT3 (HPA059914, Sigma-Aldrich), FLAG M2 (F1804, Sigma-Aldrich), MAVS E3 (sc-166583, Santa Cruz), RIG-I (sc-376845, Santa Cruz), Myc (M4439, Sigma-Aldrich). Afterwards, the membranes were washed 3x using PBST. Binding of the primary antibodies was detected using secondary antibodies (anti-rabbit IRDye 800CW (926-32211), anti-rabbit IRDye 680RD (926-68071), anti-rat IRDye 680RD (926-68076), anti-mouse IRDye 800CW (926-32210)) sourced from LI-COR. After 1.5 h incubation with 1:10,000 dilution of secondary antibodies in blocking buffer, membranes were washed 3x with PBST and fluorescent signals were detected using an Odyssey DLx scanner (LI-COR). Signal quantification was done using the Image Studio Lite software (LI-COR).

### RNP reconstitution and *IFN-β* promoter activity assay

HEK293T cells were transfected in 24-well plate format with 0.25 μg/well of pcDNA3 plasmids encoding A/WSN/33 PB2, PB1 (wild-type, T677A or D445A/D446A (PB1a) mutants), PA (wild-type, D108A or K134A mutants) and NP (wild-type or N319K mutant) and 0.25 μg/well of pPolI plasmid encoding a full-length vRNA segment or an internally truncated segment (Supplementary Table 8) under the control of the PolI promoter. NA d10A template contained a deletion of base 10 (adenosine) from the 5ʹ vRNA terminus as described in^42^. Polymerases of A/Brevig Mission/1/1918 H1N1, A/Fuijan/1/1995 H5N1, A/Vietnam/1203/2004 H5N1, A/Bilthoven/16190/1968 H3N2, and A/Netherlands/499/2017 H3N2 strains have also been used. For assessment of *IFN-β* promoter activity, 0.1 μg of a plasmid expressing firefly luciferase from the *IFN-β* promoter and 0.01 μg of a plasmid expressing *Renilla* luciferase driven by the chicken β-actin promoter with a CMV enhancer were added to the transfection mix. The transfection was performed using a 1: 2.5 DNA (μg): Lipofectamine 2000 (Invitrogen) ratio. Twenty-four hours later, the complete DMEM was aspirated, cells detached with PBS, and split into fractions for RNA isolation (1/2 cell pellet), western blotting (1/4 cell pellet) or the *IFN-β* promoter activity assay (1/4 cell pellet). For *IFN-β* promoter activity assays, 25 µl of DualGlo reagent (Promega) was added to 25 μl of cells suspended in PBS. The mixture was incubated for 10 min and the firefly luciferase readings were taken using a GloMax microplate reader (Promega) or a Synergy LX plate reader (BioTek). Next, 25 µl of Stop-Glo reagent/well was added, incubated for 10 min, and the *Renilla* luciferase readings taken. Firefly luciferase values were normalised by the *Renilla* luciferase values.

### Removal of capped or polyadenylated RNA

To remove capped RNAs, total RNA extracted from RNP reconstitution assays was treated sequentially with mRNA Decapping Enzyme (NEB) and XRN-1 (NEB) using manufacturer’s protocols. For removal of polyadenylated RNA, total RNA was treated with oligo d(T)_25_ beads from magnetic mRNA isolation kit (NEB). The resulting RNA was purified using RNA Clean & Concentrator-5 (Zymo Research).

### Primer extension assay

Primer extensions were performed essentially as described previously^56^. Briefly, total RNA was reverse-transcribed using Superscript III (Invitrogen) and ^32^P-labelled primers targeting the viral RNA of interest and the cellular 5S rRNA loading control (Supplementary Table 7) in 10 μl reactions at 50 °C for 1 h. Reactions were stopped using 10 μl formamide loading dye (90% formamide, 10 mM EDTA, 0.25% bromophenol blue, 0.25% xylene cyanol FF) and denatured at 95 °C for 5 min. cDNA products were resolved on a denaturing acrylamide gel (6% acrylamide (19:1, Bio-Rad), 7 M urea, 1x Tris-borate EDTA buffer (2 mM EDTA, 89 mM boric acid, 89 mM Tris), 0.1% TEMED, 0.06% (w/v) APS) in 1x Tris-borate EDTA running buffer for 1-1.5 h at 35 W. Afterwards, the gel was dried and the radiolabelled signals visualised using phosphorimaging plates (FujiFilm) on a Typhoon FLA 9000 scanner (GE Healthcare). Signal quantification was done using Image Studio Lite software (LI-COR).

### IAV RNA polymerase purification

Wild-type and mutant IAV RNA polymerases were purified using tandem-affinity purification (TAP) as described previously^56^ with modifications. Briefly, 4 μg of pcDNA3 plasmids expressing wild-type or mutant PB1, PA, and PB2-TAP were transfected into HEK293T cells using 60 µg PEI in Opti-MEM (Gibco). Forty-eight hours after transfection, cells were harvested in PBS, washed once with PBS, and then lysed in 1 ml lysis buffer (50 mM HEPES pH 7.5, 200 mM NaCl, 25% glycerol (Sigma), 2% Tween-20 (RPI), 1 mM β-mercaptoethanol (Bio-Rad), and 1x EDTA-free Protease inhibitor cocktail (Roche)) at 4 °C for 1 h. Next, the lysed cells were sonicated to disrupt DNA and RNA molecules, and centrifuged at 17000 x *g* for 5 min at 4 °C. The cleared lysates were bound to 50 μl IgG Sepharose beads 6 Flow (GE Healthcare) that were pre-washed 3x in binding buffer (50 mM HEPES pH 8, 200 mM NaCl, 25% glycerol, 2% Tween-20). The binding was performed under constant rotation at 4 °C. After 2 h, the beads were washed 3x with binding buffer and once with cleavage buffer (50 mM HEPES pH 7.5, 200 mM NaCl, 25% glycerol, 0.5% Tween, 1 mM DTT). Finally, the TAP-tag was removed using Tobacco etch virus (TEV) protease (Invitrogen, #12575-015) in 250 μl of cleavage buffer. Cleavage was performed at 4 °C for approximately 16 h. The beads were separated from the cleaved RNA polymerase by centrifugation at 500 *x g* for 1 min. The partially purified RNA polymerase was analysed through SDS-PAGE and silver staining using a Pierce silverstain kit (Invitrogen).

### Activity assays with the purified IAV polymerase

To measure the ability of the IAV RNA polymerase to extend a capped primer, first a synthetic 11-nt long RNA with 5ʹ diphosphate (Chemgenes; Supplementary Table 9) was capped with a radiolabelled cap-1 structure in 20 µl reactions containing 1 µM RNA, 0.25 µM [α-^32^P]GTP (3000 Ci/mmole, Perking-Elmer), 0.8 mM S-adenosylmethionine, 0.5 U/µl *Vaccinia virus* capping enzyme (NEB), and 2.5 U/µl 2ʹ-O-methyltransferase (NEB) at 37 °C for 1 h. Next, the capped RNA was purified using an oligonucleotide cleanup kit (Zymo Research) and eluted in 30 µl water. To test the transcriptional activity of the IAV RNA polymerase, we set up 4-μl reactions containing 5 mM MgCl_2_, 1 mM DTT, 2 U/μl RNase inhibitor (APExBIO), 0.5 μM of NA71 or NA71-U RNA template (Supplementary Table 9), 500 μM ATP, 500 μM CTP, 500 μM GTP, ∼1500 cpm capped RNA primer, 12.5% glycerol, 1% Tween-20, 100 mM NaCl, 25 mM HEPES pH 8, 50 µM baloxavir (AstaTech), and 5 ng/µl RNA polymerase. Reactions were incubated at 30 °C for 1 h and analysed with 12% denaturing PAGE and autoradiography.

For the RNase H analysis, transcription reactions were performed as described above and stopped by boiling at 95 °C for 2 min. Next, 1 μM oligo d(T)_20_ and 2.5 U RNase H (NEB) were added and the samples incubated at 37 °C for 30 min. The RNase-treated reactions were analysed with 12% denaturing PAGE and autoradiography.

To measure the cap snatching activity of the RNA polymerase, we capped a synthetic 20-nt long RNA with a 5ʹ-diphosphate (Chemgenes, Supplementary Table 9) with a radiolabelled cap-1 structure in 20 µl reactions as described above for the 11-nt long RNA. The capped 20-nt long RNA was purified using an oligonucleotide cleanup kit (Zymo Research) and eluted in 30 µl RNase free water. To test the cleavage activity, 4-µl reactions were performed as described above with the modification that NA71 template was replaced with the model vRNA promoter (vRNA_5p and vRNA_3p, Supplementary Table 9), but NTPs and baloxavir were omitted and replaced with RNase-free water. Reactions were analysed through 12% denaturing PAGE and autoradiography.

### Polymerase-vRNA binding assay

Recombinant polymerases were purified using a TAP-tag on the C-terminus of the PB2 subunit. Briefly, HEK293T cells were transfected with 4 μg of pcDNA3 plasmids encoding PB1 (wild-type, T677A or R238A/R672A), TAP-tagged PB2, PA and a pPolI plasmid encoding full-length segment 6 in a 10 cm dish using Polyethylenimine (PEI, Sigma) transfection reagent at a 1: 3 (DNA (μg): PEI (μl)) ratio. Forty-eight hours after transfection cells were lysed with cold lysis buffer (50 mM HEPES (pH 8.0), 200 mM NaCl, 25% glycerol, 0.5% NP-40, 0.007% β-mercaptoethanol, 1x cOmplete protease inhibitor cocktail (Roche)) on ice. Debris were removed by centrifugation at 13,000 g and 4 °C for 5 min. Next, the NaCl concentration of the supernatant was returned to 150 mM. The supernatant was incubated with 50 µl of IgG Sepharose beads (GE Healthcare) (pre-washed 5x with binding buffer (10 mM HEPES (pH 8.0), 150 mM NaCl, 10% glycerol, 0.1% NP-40)) at 4 °C for 2 h. Next, the IgG Sepharose beads were washed 3x with binding buffer and RNA and protein samples were collected from the total lysate and the bead-bound fractions. The RNA samples were analysed using primer extension with NA 1280 and 5S 100 primers (Supplementary Table 7).

### RT-PCR for detection of viral genome segments and DelVGs

Reverse transcription reactions were done using total RNA isolated from infected cells or virus-containing supernatants, Tuni-12 and Tuni-13 primers^63^ (Supplementary Table 7), and SuperScript III (Invitrogen) at 50 °C for 1 h. PCR reactions were done using Tuni-12 and Tuni-13 primers and *Pfu* DNA polymerase (Promega). PCR products were resolved on a 1.5% 0.5x TAE agarose gel.

### RT-PCR for detection of mvRNA

RNA isolated from infection samples was fractionated by length using an RNA Clean & Concentrator kit (Zymo Research). The small (<200 nt) RNA fraction was used for reverse transcription with Lv3aa and Lv3ga primers and SuperScript III (Invitrogen) at 37 °C for 30 min as described previously^26^. Second strand synthesis was done using the Lv5 primer and *Pfu* DNA polymerase (Promega) at 95 °C for 2 min, 47 °C for 10 min and 72 °C for 3 min. Next, samples were treated with Exonuclease VII (NEB) at 37 °C for 30 min and 95 °C for 10 min. PCR was done with P7 and P5_IRdye800 (conjugated to an IRdye800 fluorophore) primers and *Pfu* DNA polymerase. Thermocycling conditions included: i) initial denaturation at 95 °C for 2 min; ii) 24-28 cycles of denaturation at 95 °C for 30 s, annealing at 59 °C for 30 s and extension at 72 °C for 30 s; iii) final extension at 72 °C for 5 min. RT-PCR for 5S rRNA was done as described above (without a specific second strand synthesis) using the 5S 100 primer for reverse transcription and 5S_Fw and 5S100_Rev_ATTO647N (conjugated to an ATTO647N fluorophore) primers for PCR. Primers used here are listed in Supplementary Table 7. Five μl of 6x Orange G loading dye (6x concentration: 10 mM Tris (pH 7.5), 50% glycerol, 1 mM EDTA, Orange G) was added and samples were run on a 6% acrylamide TBE gel. RT-PCR products were visualised using 700 nm (for 5S rRNA) or 800 nm (for mvRNA) wavelengths on an Odyssey DLx scanner (LI-COR). Gels were subsequently stained with SYBR Gold (Invitrogen) to visualise the ladder using a G:BOX XX6 gel imager (Syngene).

### Immunoprecipitation of Myc-tagged RIG-I

For immunoprecipitation assays with full-length IAV segments, HEK293T cells were transfected with 3 µg of each of the following plasmids: pcDNA3-NP, pcDNA3-PB2, pcDNA3-PA, pcDNA3-PB1 (wild-type or T677A mutant), pcDNA-Myc-RIG-I (wild-type or K861A/K858A/K851A mutant) and pPolI plasmid encoding a full-length NA or NP segment. The transfection was done using 1: 2.5 DNA (μg): Lipofectamine 2000 (Invitrogen) ratio in 10 cm dishes. Forty-eight hours later, cells were detached with 10 ml of cold PBS, pelleted at 1,000 rpm for 5 min at 4 °C and lysed in 600 µl of cold lysis buffer (50 mM Tris-HCl (pH 8.0), 5% glycerol, 0.5% NP-40, 200 mM NaCl, 1 mM EDTA, 1 mM DTT, cOmplete protease inhibitor cocktail (Roche)) at 4 °C for 1 h. The cell debris were removed by pelleting at 13,000 rpm, 4°C for 5 min and 20% of the sample was taken for ‘total’ RNA and protein analyses. The remaining lysate was incubated with 1.5 µl of anti-Myc antibody (#M4439, Sigma-Aldrich) at 4°C for 2 h under rotation. Twenty-five µl of Protein G Mag Sepharose Xtra beads (GE Healthcare) per condition was washed 3x with 1 ml of wash buffer (10 mM Tris-HCl (pH 8.0), 150 mM NaCl, 0.1% NP-40, 1 mM EDTA). Then lysates were added to the beads and incubated overnight at 4 °C under rotation. Next, beads were washed 3x with wash buffer at 4 °C for 5 min under rotation. Finally, two thirds of the bead suspension was used for RNA extraction and one third for protein analysis. RNA samples were analysed by primer extension with NA160 and NA1280 or NP 149- and NP149+ primers (Supplementary Table 7).

For immunoprecipitations with the NA196 template, 10 cm dishes with HEK293T cells were transfected with 4 µg of each of the following plasmids: pcDNA3-NP, pcDNA3-PB2, pcDNA3-PA, pcDNA3-PB1 (wild-type or T677A mutant), pcDNA-Myc-RIG-I (wild-type, K861A/K858A/K851A, K270A mutant) or empty pcDNA3 plasmid and pPolI plasmid encoding the NA196 template. Transfections were done using a 1: 5 (DNA (μg): PEI (μl)) ratio. Forty-eight hours after transfection, cells were washed with cold PBS and lysed with 600 µl of cold lysis buffer (10 mM Tris-HCl pH 7.5, 150 mM NaCl, 0.5% NP-40, 1x cOmplete protease inhibitor cocktail (Roche)) on ice for 1 h. Cell debris was removed through centrifugation at 17,000 rpm, 4 °C for 20 min. Next, 900 µl of dilution buffer (10 mM Tris-HCl (pH 7.5), 150 mM NaCl, cOmplete protease inhibitor cocktail (Roche)) was added to the supernatants. Ten percent of the supernatants was saved for ‘total’ RNA and protein analyses. Twenty-five µl of ChromoTek Myc-Trap® Agarose beads (Proteintech) (pre-wasehed 3x with wash buffer (10 mM Tris-HCl (pH 7.5), 200 mM NaCl, 0.05% NP-40)) were added to the remaining supernatant and incubated at 4 °C for 1 h under rotation. Next, the beads were washed 3x with wash buffer and two-thirds of the bead suspension was used for RNA extraction and one third for protein analysis. RNA samples were analysed by primer extension with the NA 5ʹ or NA-2 primer (Supplementary Table 7). RIG-I pull-down with the full-length NA temple for TSO-based RT-PCR was done the same way, except immunoprecipitation was done 24 h after transfection.

For RIG-I immunoprecipitation in infection, HEK293T cells were transfected with 15 µg of pcDNA-empty, pcDNA encoding wild-type RIG-I or a K861A/K858A/K851A mutant RIG-I at a 1:5 (DNA (µg): PEI (µl)) ratio. Twenty-four hours after transfection cells were infected with wild-type or T677A mutant A/WSN/33 viruses at an MOI equivalent to an MOI of 0.01 of T677A virus stock. The inoculum volume of the wild-type virus was adjusted to the T677A mutant virus based on qPCR results for PB1 vRNA to normalise the incoming RNA levels. Immunoprecipitation was done 24 h after infection as described for NA196 template. RNA levels were analysed with TSO-based RT-PCR.

### TSO-based RT-PCR and gel electrophoresis

For optimisation of the TSO-based RT-PCR assay, 0.625 ng of *in vitro* transcribed vRNA, cRNA, ccRNA, ccRNA-3ʹ, mRNA or combinations thereof were mixed with 10 ng of total HEK293T RNA and subsequently used as input for the reactions. For the TSO-based RT-PCR analysis of tissue culture infection samples, RNA was extracted from A549 cells infected with an MOI of 0.3 of wild-type or T677A A/WSN/33 virus for 24 h. For analysis of mouse infection samples, RNA was isolated from mouse lungs 1 day after infection with 10^5^ PFU of A/Vietnam/1203/04 (H5N1). Next, RNA was extracted, and 2 µg of total RNA was either treated or mock-treated with oligo d(T)_25_ beads from the magnetic mRNA isolation kit (NEB). RNA was subsequently purified using an RNA Clean & Concentrator-5 kit (Zymo Research). TSO-based RT was done with equal volumes (containing 100-200 ng) of RNA. To this end, RNA was first denatured in the presence of dNTPs (1mM final concentration) and either the Tuni-13 LNA3, or the oligo d(T)_20_ primer (1 µM final concentration) in 3 µl volume at 70 °C for 5 min. Denatured RNAs were immediately placed on ice for 5 min. Next, 2 µl of enzyme mix containing template switching RT buffer (NEB), template switching RT enzyme mix (NEB), and the TSO (3.75 µM final concentration) were added to the RNA mix and incubated at 42 °C for 90 min. Reactions were terminated by incubation at 85 °C for 5 min and subsequently cooled to 4 °C. The unused primers were digested with Thermolabile Exonuclease I (NEB) and reactions were diluted 2x with water. One µl of diluted RT reaction was used for PCR amplification with Q5 High-Fidelity DNA polymerase (NEB) using TSO-specific forward primer and a gene-specific reverse primer. Primers and oligos used here are listed in Supplementary Table 7. Thermocycling conditions included: i) initial denaturation at 98 °C for 30 s; ii) 25-29 cycles of denaturation at 98 °C for 10 s, annealing at 50 °C for 15 s and extension at 72 °C for 10 s; iii) final extension at 72 °C for 2 min. Note that due to abundance of mRNA in infection, lower number of PCR amplification cycles was used for mRNA than for cRNA/ccRNA to avoid oversaturation. The PCR products were resolved on 8-10% TBE acrylamide gel and visualised using SYBR Gold (Invitrogen) and G:BOX XX6 gel imager (Syngene).

### Quantitative PCR

To determine the efficiency of the mRNA depletion with oligo d(T)_25_ beads, RNA from TSO-based RT reactions with either the Tuni-13 LNA3, oligo d(T)_20_ or 18S rRNA reverse primer was treated with Thermolabile Exonuclease I (NEB) and used for qPCR. qPCR reactions were done using NA-specific or 18S rRNA-specific primers and qRT-PCR Brilliant III SYBR Master Mix with ROX (Agilent). To normalise the viral inoculum volumes for RIG-I immunoprecipitation experiments, RT was done using Tuni-12 and 18S rRNA Rev primers qPCR was done with PB1- and 18S rRNA-specific primers (Supplementary Table 7).

### Sample preparation for sequencing using Oxford Nanopore Technology (ONT)

Spike RNA for sequencing was prepared using *in vitro* transcription and was based on the segment eight cRNA sequence that contained a specific, not natural capped primer-like sequence at the 5ʹ terminus and a barcode at the 3ʹ terminus to control for intermolecular ligation. The spike sequence also contained two additional barcodes adjacent to the RT and PCR primer binding sites, that were used to estimate the number of template switching events during RT (Supplementary Table 9, Supplementary Fig. 6). The DNA template for *in vitro* transcription reaction was synthesised by gBlocks^TM^ (IDT) and PCR-amplified using Q5 High-Fidelity DNA polymerase (NEB) and NS spike T7 PCR Fw and Rev primers (Supplementary Table 7). *In vitro* transcription reactions were done using a MEGAshortscript™ T7 Transcription Kit (ThermoFisher) and the resulting product RNA was purified using an RNA Clean & Concentrator-5 kit (Zymo Research). Spike RNA was then treated with RppH (NEB) and NEBuffer2 (NEB) according to for one hour at 37 °C and purified using RNA Clean & Concentrator-5 (Zymo Research).

To prepare samples for sequencing HEK293-luc cells were infected with wild-type or T677A mutant viruses at an MOI of 3 for one hour at 4 °C (in a 24-well plate). After washing with PBS and replacing infection media, cells were transferred to 37 °C for 6 hours. Two µg of total RNA from infections were treated with 25 μl Oligo d(T)_25_ magnetic beads (NEB). The RNA was then decapped using mRNA decapping enzyme (NEB) and 5ʹ pyrophosphates were removed using RppH (NEB) and NEBuffer 2 (NEB) through a 1 h incubation at 37°C. One μg of treated RNA from the infections was then circularised in the presence of 0.02 ng of RppH-treated spike RNA using T4 ssRNA Ligase (NEB) at 10% PEG 8000 at 16 °C overnight. RNA was cleaned up using RNA Clean & Concentrator-5 (Zymo Research) after each step. Circularised RNA was then reverse transcribed using SuperScript III (Invitrogen) and NS seq Rev primer, and RNA was digested using RNase H (NEB). cDNA was then PCR-amplified using Q5 High-Fidelity DNA polymerase (NEB) and NS seq Fw and NS seq Rev primers for 30 cycles (Supplementary Table 7). The PCR products were resolved on a 1% TAE agarose gel and bands of around 850 bp size were excised and purified with Monarch PCR and DNA Cleanup Kit (NEB) and RNAClean XP beads (Beckman Coulter). DNA concentration and purity were assessed with Qubit and nanodrop respectively.

Samples were submitted for Plasmidsaurus custom sequencing. The library was prepared using ONT’s kit SQK-NBD114.96. Sequencing was performed on PromethION P24 instrument with R10.4.1 flow cell, minimum Qscore 10, minimum read length 200 bp. Basecalling was performed with Dorado v4.3.0 on super-accurate mode, and adapters and barcodes were trimmed by MinKNOW software.

### Computational analysis of the ONT sequencing

Sequence analysis was done using SeqKit^64^ as shown in Supplementary Fig. 7 and described in the Supplementary Note. Briefly, raw Oxford Nanopore sequencing reads were size-selected, with reads shorter than 800 nt or longer than 1000 nt excluded from the analysis. Next, reads were binned into ccRNA and cRNA pools based on the presence or absence (respectively) of extensions in the junction between the cRNA 3ʹ and 5ʹ termini. Reads with a complete 5ʹ cRNA terminus and those lacking the first adenosine from the 5ʹ cRNA terminus were counted as cRNAs and quantified. The sequence extensions between cRNA 3ʹ and 5ʹ in the ccRNA pool were isolated and sequences containing multiple cRNA 3ʹ and 5ʹ termini were filtered out. The number of spike RNA 5ʹ reads among the sequence extensions was used to estimate the intermolecular ligation rate. The spike RNA 5ʹ sequence was subsequently filtered out from the ccRNA pool. Molecules with a capped primer length of 8-30 nt between cRNA 3ʹ and 5ʹ were considered ccRNAs. The cRNA and ccRNA pools were also used to find reads with extended internal A-stretches.

The spike to viral ratio was calculated using reads containing spike 3ʹ and 5ʹ termini and those containing the viral cRNA 3ʹ and 5ʹ termini. The rate of template switching was estimated using spike barcodes adjacent to the RT and PCR primer biding sites.

All custom analysis scripts were developed with the assistance of Google Gemini v3.5. Scripts and output were inspected manually for errors. All data visualizations pertaining to ONT analysis were generated using Python version 3.13 and the Matplotlib library^65^.

### *In vitro* transcription

DNA templates for the *in vitro* transcription reactions were prepared by PCR using the primers listed in Supplementary Table 7, Q5 High-Fidelity DNA polymerase (NEB), and the pPolI-NA196 plasmid as a template. The DNA template for the svRNA *in vitro* transcription was prepared by annealing svRNA forward and reverse oligos (Supplementary Table 7) using a stepwise protocol (90 °C for 1 min, ramping down to 25 °C over 1h, and holding at 4 °C). *In vitro* transcriptions were done using MEGAshortscript™ T7 Transcription Kit (ThermoFisher). The products were resolved on a denaturing acrylamide gel (6% or 15% Acrylamide (19:1, Bio-Rad) (for longer RNA or svRNA respectively), 7M urea, 1x Tris-borate EDTA buffer (2 mM EDTA, 89 mM boric acid, 89 mM Tris), 0.1% TEMED, 0.06% (w/v) APS) in 1x Tris-borate EDTA running buffer. The products were gel-purified by freezing gel slices on dry ice, crushing the slices, and incubating the slices with elution buffer (20 mM Tris-HCl pH 7.5, 0.25 M sodium acetate, 1 mM EDTA, and 0.25% SDS) (for longer RNAs) or water (for svRNA) at RT overnight, under constant shaking. The RNA was subsequently purified using RNA Clean & Concentrator-5 (Zymo Research), with the modification of increasing the ethanol concentration during column binding to 66% for the svRNA purification. Part of the cRNA sample was dephosphorylated using Antarctic phosphatase (NEB). The ccRNA, mRNA and ccRNA-3′ transcripts were capped and methylated using Vaccinia Capping Enzyme (NEB) and cap 2’-O-methyltransferase (NEB). Afterwards, ccRNA, mRNA and ccRNA-3′ were treated sequentially with Antarctic phosphatase (NEB), PNK (NEB) and XRN-1 (NEB) to remove uncapped RNA. The RNA was purified using RNA Clean & Concentrator-5 (Zymo Research) in between treatments. The quality of the final RNA preparations was assessed using denaturing acrylamide gel electrophoresis. Sequences of expected *in vitro* transcription products are listed in Supplementary Table 8.

### RNA transfection

The *in vitro* transcribed RNAs were transfected alone or in combination with svRNA or pPolI plasmids encoding segment 6- or segment 5 -based vRNAs into HEK293-luc or HEK293-luc *RIG-I* -/- cells together with *Renilla* luciferase-encoding plasmid and RNAse inhibitor (APExBIO) using Lipofectamine 2000 (Invitrogen) reagent. In all transfections, the *in vitro* transcribed RNA was diluted in total HEK293T RNA, which was also used as a negative control. The MAVS-FLAG encoding plasmid was used as a positive control as described previously^54^. Firefly luciferase and *Renilla* luciferase readings were taken using Dual-Glo® Luciferase Assay System (Promega) 20 h post-transfection.

### HEK293-luc wild-type, *RIG-I* -/- and *MAVS* -/- cell line validation

To validate the firefly luciferase reporter activity under the *IFN-β* promoter, HEK293-luc wild-type, *RIG-I* -/- and *MAVS* -/- cells were transfected with 50 ng/well MAVS-FLAG expressing plasmid or an empty control plasmid along with *Renilla* luciferase reporter plasmid (in 96-well plate) using Lipofectamine 2000 (Invitrogen) reagent. Firefly luciferase and *Renilla* luciferase readings were taken using Dual-Glo® Luciferase Assay System (Promega) 24 h post-transfection. To test activity of different innate immune signalling pathways, HEK293-luc wild-type, *RIG-I* -/- and *MAVS* -/- cells were transfected with 20 ng/well of HMW poly(I:C) (Invivogen), LMW poly(I:C) (Invivogen), *in vitro* transcribed NA196 vRNA, HEK293T RNA (mock) or 100 ng/well HSV-1 60-mer DNA (in 96-well plate) along with 20 ng *Renilla* luciferase reporter plasmid and 40 U of RNAse inhibitor (APExBIO) using Lipofectamine 2000 (Invitrogen) reagent. Prior to transfection HSV-1 60-mer DNA forward and reverse oligos (Supplementary Table 7) were annealed using a stepwise protocol (90 °C for 1 min, ramping down to 25 °C over 1h, and holding at 4 °C). Firefly luciferase and *Renilla* luciferase readings were taken using Dual-Glo® Luciferase Assay System (Promega) 20 h post-transfection.

### RIG-I activation assay

RIG-I was purified as described previously^66^. For ATPase assays, 0.5 µM RIG-I was incubated with 0.1 µM [γ-^32^P]ATP and 10 ng of template RNA. Activity assays were performed in a buffer containing 50 mM HEPES pH 8.0, 150 mM NaCl, 2 mM MgCl_2_, and 5 mM DTT, and quenched using 1M formic acid/50 mM EDTA. [γ-^32^P]ATP and ^32^P_i_ were resolved using glass-backed PEI-cellulose TLC plates (Sigma-Aldrich) in 0.4 M KH_2_PO_4_ (pH 3.4). After wrapping the TLC plates in plastic foil, the radioactive signals were detected using BAS-MS phosphorimaging plates (FujiFilm) through a 1 h exposure and visualised using a Typhoon FLA 9000 scanner (GE Healthcare). Densitometry analysis was performed using ImageJ.

### Statistical analysis

In all figures, error bars indicate standard deviation with the number of biological repeats indicated in figures or figure legends. Statistical analysis was done using GraphPad Prism Software. Details on the tests used are indicated in each figure legend.

## Acknowledgements

The authors would like to thank members of the te Velthuis lab for helpful discussions, and Jennifer Miller and Jean Arly Volmar in the Princeton Genomics Core Facility for assistance with the NGS of the virus passaging experiment.

## Author contributions

EE, EP, JF, KB, AYJ, and AJWtV performed biochemical experiments. EE, WW, MM, and ABR performed sequencing. HF and JWP generated mutant RIG-I. BS and JM purified RIG-I. EE, MSO, ABR, and AJWtV analysed data. EE and AJWtV wrote the manuscript. All authors approved the manuscript.

## Data availability

NGS data from virus passaging and ONT sequencing have been deposited in the NCBI database under the accession code PRJNA1177455. Raw image, the ONT sequencing analysis script, and numerical data are available via Zenodo at https://doi.org/10.5281/zenodo.15102697. The script is also available via Github through https://github.com/ajwtevelthuis/ccRNA-analysis. Plasmids are available via https://www.addgene.org/AJ_te_Velthuis/.

