## Supplemental Tables for "Aberrant influenza virus transcription termination generates capped cRNAs that activate RIG-I"

**Supplementary Table 1. Weights and lung viral titres of mice infected**

| mouse | inoculation | Weight day 0 |
|---|---|---|
| Mock | PBS | 20.20 g (100%) |
| Infected 1 | A/Vietnam/1203/04 (H5N1) | 19.90 g (100%) |
| Infected 2 | A/Vietnam/1203/04 (H5N1) | 18.60 g (100%) |
| Infected 3 | A/Vietnam/1203/04 (H5N1) | 20.30 g (100%) |

**with A/Vietnam/1203/04 (H5N1)**

| Weight day 1 | virus titre (pfu/ml) |
|---|---|
| 19.75 g (97.77%) | <100 |
| 18.75 g (94.22%) | 160000 |
| 17.83 g (95.86%) | 270000 |
| 19.14 g (94.29%) | 210000 |

**Supplementary Table 2. Raw read numbers and calculations for te[illegible]**

| sample | Raw reads | 800-1000 nt |
|---|---|---|
| wild-type replicate 1 | 778457 | 303443 |
| wild-type replicate 2 | 628848 | 297846 |
| wild-type replicate 3 | 665458 | 289212 |
| T677A replicate 1 | 656643 | 281665 |
| T677A replicate 2 | 735013 | 300707 |
| T677A replicate 3 | 627509 | 298820 |

**mplate switching**

| cRNA Fw+cRNA Rev (same) | spike Fw+spike Rev (same) |
|---:|---:|
| 92527 | 100375 |
| 38572 | 154062 |
| 42568 | 159337 |
| 133852 | 62127 |
| 101731 | 82307 |
| 64668 | 138724 |

| cRNA Fw+spike Rev (switched) | spike Fw+cRNA Rev (switched) |
|---|---|
| 38203 | 23535 |
| 26664 | 16799 |
| 31588 | 18084 |
| 36313 | 18756 |
| 35300 | 18738 |
| 36798 | 20452 |

| contains both same and switched | lacks complete pair | % template switching |
|---|---|---|
| 210 | 48593 | 24.25 |
| 124 | 45444 | 18.41 |
| 141 | 46128 | 19.74 |
| 190 | 49469 | 21.94 |
| 155 | 50981 | 22.70 |
| 133 | 38045 | 21.96 |

**Supplementary Table 3. Raw read numbers and calculation**

| sample | Raw reads | 800-1000 nt |
|---|---|---|
| wild-type replicate 1 | 778457 | 303443 |
| wild-type replicate 2 | 656643 | 281665 |
| wild-type replicate 3 | 628848 | 297846 |
| T677A replicate 1 | 735013 | 300707 |
| T677A replicate 2 | 665458 | 289212 |
| T677A replicate 3 | 627509 | 298820 |

**s for capped primer lengths, spike to viral RNA ratio and intermolecular ligation**

| cRNA 3' and 5' (one pair, same strand) | spike 3' and 5' (one pair, same strand) |
|---|---|
| 102298 | 127168 |
| 49534 | 162854 |
| 56316 | 167860 |
| 87467 | 87668 |
| 61073 | 102004 |
| 47241 | 153692 |

| spike/viral ratio | cRNA (missing first A at 5') | cRNA (complete 5') | cRNA total |
|---|---|---|---|
| **1.24** | 1364 | 98436 | 99800 |
| **3.29** | 522 | 47861 | 48383 |
| **2.98** | 695 | 54236 | 54931 |
| **1.00** | 3877 | 65916 | 69793 |
| **1.67** | 2349 | 46708 | 49057 |
| **3.25** | 1968 | 35202 | 37170 |

| cRNA missing 1st A at 5' (% cRNA total) |
|---:|
| 1.37 |
| 1.08 |
| 1.27 |
| 5.55 |
| 4.79 |
| 5.29 |

| reads with extension between 3' and 5' (no multimappers, with spike) |
| ---: |
| 2621 |
| 1179 |
| 1424 |
| 17825 |
| 12056 |
| 10103 |

| reads with extension between cRNA 3' and 5' containing 5' spike |
|---:|
| 76 |
| 25 |
| 23 |
| 63 |
| 25 |
| 16 |

| reads with extension between 3' and 5' (no multimappers, no spike) |
|---:|
| 2545 |
| 1154 |
| 1401 |
| 17762 |
| 12031 |
| 10087 |

| % intermolecular ligation | ccRNA (8-30 nt cap seq) |
|---:|---:|
| **0.074204** | 1341 |
| **0.050442** | 539 |
| **0.040813** | 728 |
| **0.071903** | 16407 |
| **0.040908** | 11086 |
| **0.033846** | 9357 |

| ccRNA (8-30 nt cap seq) (% of total cRNA + reads with extension between 5' and 3 cRNA ( |
|---:|
| 1.31 |
| 1.09 |
| 1.29 |
| 18.73 |
| 18.14 |
| 19.79 |

| ccRNA (8-30 nt cap seq) (% of total cRNA + reads with extension between 5' and 3 cRNA ([illegible] |
| ---: |
| 1.31 |
| 1.09 |
| 1.29 |
| 18.74 |
| 18.15 |
| 19.80 |

spike))

**Supplementary Table 4. Raw read numbers a**

| sample | cRNAs total |
|---|---|
| wild-type replicate 1 | 99800 |
| wild-type replicate 2 | 48383 |
| wild-type replicate 3 | 54931 |
| T677A replicate 1 | 69793 |
| T677A replicate 2 | 49057 |
| T677A replicate 3 | 37169 |

**nd calculations for the A-stretch length**

| cRNAs with 6 As in A-stretch | cRNAs with >6 As in A-stretch |
|---|---|
| 91216 | 991 |
| 44578 | 494 |
| 50399 | 592 |
| 63857 | 799 |
| 44839 | 482 |
| 34337 | 354 |

| cRNAs with < 6 As or other nt in A-stretch | cRNAs that do not posess a valid pattern |
|---:|---:|
| 1512 | 6081 |
| 682 | 2629 |
| 800 | 3140 |
| 961 | 4176 |
| 713 | 3023 |
| 513 | 1965 |

| total cRNAs with 6 As or more in A-stretch | % cRNAs with extended A-stretch |
|---|---|
| 92207 | **1.07476** |
| 45072 | **1.09602** |
| 50991 | **1.16099** |
| 64656 | **1.23577** |
| 45321 | **1.06352** |
| 34691 | **1.02044** |

| ccRNAs total | ccRNAs (8-30 caps) with 6 As in A-stretch |
|---|---|
| 1341 | 1145 |
| 539 | 477 |
| 728 | 644 |
| 16407 | 13757 |
| 11086 | 9096 |
| 9357 | 7762 |

| ccRNAs (8-30 caps) with >6 As in A-stretch |
|---:|
| 88 |
| 24 |
| 40 |
| 1503 |
| 1205 |
| 1050 |

| **ccRNAs with < 6 As or other nt in A-stretch** |
|---:|
| 24 |
| 12 |
| 12 |
| 235 |
| 151 |
| 116 |

| ccRNAs that do not posess a valid pattern |
| ---: |
| 84 |
| 26 |
| 32 |
| 912 |
| 634 |
| 429 |

| total ccRNAs with 6 As or more in A-stretch |
|---|
| 1233 |
| 501 |
| 684 |
| 15260 |
| 10301 |
| 8812 |

| % ccRNAs (8-30 caps) with extended A-stretch |
|---:|
| 7.14 |
| 4.79 |
| 5.85 |
| 9.85 |
| 11.70 |
| 11.92 |

**Supplementary Table 5. Variants identifie**

| Segment | position | type |
| --- | --- | --- |
| 3 (PA) | 1711 | snp |
| 6 (NA) | 66 | snp |
| 5 (NP) | 143 | snp |
| 6 (NA) | 1368 | snp |
| 6 (NA) | 1103 | snp |
| 6 (NA) | 1325 | snp |
| 6 (NA) | 1330 | snp |
| 3 (PA) | 244 | del |
| 3 (PA) | 166 | del |
| 3 (PA) | 51 | del |
| 3 (PA) | 1943 | del |
| 1 (PB2) | 2300 | snp |
| 2 (PB1) | 1882 | del |

**ed in the stock and passages of the wild-type A/WSN/33 virus**

| reference | allele | Amino acid/ORF change |
|---|---|---|
| T | C | silent |
| T | C | V16A |
| T | C | I33T |
| C | G | T450S |
| A | G | N362D |
| A | G | T436A |
| A | G | silent |
| CACAGATTTGAAATA | CA | frameshift (81 aa product) |
| TATTCA | TA | stop (47 aa product) |
| CAATC | CATC | frameshift (11 aa product) |
| TATTGGCAAAGTCGGTATTCAACAGCTT | TA | del 641-651 |
| T | C | I760T |
| TACCAGGGGCGTTTATGCA | TA | stop (619 aa product) |

| Variant frequency (%) in virus stock | Variant frequency (%) in P7 (rep 1) |
|---|---|
| 99 | 99 |
| 82 | |
| 74 | 36 |
| | 100 |
| | 82 |
| | 78 |
| | 77 |
| | 30 |
| | 26 |
| | 26 |
| | 15 |
| | 14 |

| Variant frequency (%) in P7 (rep 2) | Variant frequency (%) in P7 (rep 3) |
|---|---|
| 99 | 99 |
| 35 | 27 |
| 100 | 100 |
| 77 | 82 |
| 78 | 71 |
| 78 | 71 |
| 14 | |

**Supplementary Table 6. Variants identified in the stock and passa**

| Segment | position | type | reference | allele |
|---|---|---|---|---|
| 2 (PB1) | 2053 | snp | A | G |
| 5 (NP) | 1002 | snp | T | G |
| 2 (PB1) | 1650 | snp | A | T |
| 2 (PB1) | 776 | snp | T | G |
| 5 (NP) | 1175 | snp | G | A |
| 4 (HA) | 1583 | snp | T | A |
| 1 (PB2) | 123 | snp | G | A |
| 5 (NP) | 37 | snp | A | G |
| 2 (PB1) | 516 | snp | A | G |
| 4 (HA) | 1136 | snp | T | C |
| 4 (HA) | 1160 | snp | C | A |
| 8 (NS) | 553 | snp | A | T |
| 5 (NP) | 1450 | snp | G | A |
| 2 (PB1) | 2226 | snp | G | A |
| 4 (HA) | 116 | snp | C | T |
| 1 (PB2) | 2226 | snp | G | A |
| 4 (HA) | 1660 | snp | T | A |
| 5 (NP) | 711 | snp | G | A |
| 3 (PA) | 243 | del | GCAC | GC |
| 3 (PA) | 115 | del | GAAACAAACAAAT | GAAACAAAT |

**ages of the T677A mutant A/WSN/33 virus**

| Amino acid/ORF change | Variant frequency (%) in virus stock |
|---|---|
| T677A | 100 |
| N319K | |
| silent | |
| F251C | |
| S377N | |
| D517E | |
| silent | |
| non-coding region | |
| I164M | |
| silent | |
| silent | |
| N176I (NS1) M19L (NEP) | |
| E469K | |
| silent | |
| silent | |
| silent | |
| stop (542 aa product) | |
| M222I | |
| frameshift (75 aa product) | |
| frameshift (59 aa product) | |

| Variant frequency (%) in P5 (rep 1) | Variant frequency (%) in P7 (rep 1) |
|---:|---:|
| 99 | 100 |
| 67 | 80 |
| 34 | 73 |
| 33 | 71 |
| 19 | 14 |
| 18 | 21 |
| 18 | 21 |
| 16 | 12 |
| 14 | 11 |
| 10 | 7 |

| Variant frequency (%) in P3 (rep 2) | Variant frequency (%) in P5 (rep 2) |
|---|---|
| 13 | 9 |
| 24 | 47 |
| 58 | 60 |
| 23 | 22 |
| 15 | 16 |
| 15 | 10 |
| 15 | 12 |
| 11 | 9 |
| 10 | 11 |

| Variant frequency (%) in P4 (rep 3) | Variant frequency (%) in P5 (rep 3) |
|---|---|
| 14 | 8 |
| 22 | 45 |
| 40 | 35 |
| 13 | 12 |
| 11 | 12 |
| 14 | 13 |
| 9 | 12 |
| 10 | 11 |
| 9 | 11 |
| 9 | 24 |
| 7 | 22 |

**Supplementary Table 7. Primers, oligos and gBlock DNA used in this study**

| Experiment | Primer/oligo/gBlock DNA name |
|---|---|
| Site-directed mutagenesis | PB1 R670A Fw |
| Site-directed mutagenesis | PB1 R670A Rev |
| Site-directed mutagenesis | PB1 N671A Fw |
| Site-directed mutagenesis | PB1 N671A Rev |
| Site-directed mutagenesis | PB1 S673A Fw |
| Site-directed mutagenesis | PB1 S673A Rev |
| Site-directed mutagenesis | PB1 N676A Fw |
| Site-directed mutagenesis | PB1 N676A Rev |
| Site-directed mutagenesis | PB1 T677A Fw |
| Site-directed mutagenesis | PB1 T677A Rev |
| Site-directed mutagenesis | PB1 S678A Fw |
| Site-directed mutagenesis | PB1 S678A Rev |
| Site-directed mutagenesis | NP N319K Fw |
| Site-directed mutagenesis | NP N319K Rev |
| Site-directed mutagenesis | RIG-I K270A Fw |
| Site-directed mutagenesis | RIG-I K270A Rev |
| Site-directed mutagenesis | RIG-I K861A/K858A/K851A |
| Site-directed mutagenesis | RIG-I K861A/K858A/K851A |
| Primer extension | 5S 100 |
| Primer extension | NP 149- |
| Primer extension | NP 149+ |
| Primer extension | NA 1280 |
| Primer extension | NA 160 |
| Primer extension | PB1 vRNA |
| Primer extension | PB1 c/mRNA |
| Primer extension | HA vRNA |
| Primer extension | HA c/mRNA |
| Primer extension | M vRNA |
| Primer extension | M c/mRNA |

| | |
|---|---|
| Primer extension | NA 5′ |
| Primer extension | NA-2 |
| Primer extension | NA c/mRNA mini |
| Primer extension | NP 5′ |
| Primer extension | GC- |
| Primer extension | NP c/mRNA mini |
| mvRNA and 5S rRNA RT-PCR | Lv3aa |
| mvRNA and 5S rRNA RT-PCR | Lv3ga |
| mvRNA and 5S rRNA RT-PCR | Lv5 |
| mvRNA and 5S rRNA RT-PCR | P7 |
| mvRNA and 5S rRNA RT-PCR | P5_IRdye800 |
| mvRNA and 5S rRNA RT-PCR | 5S 100 |
| mvRNA and 5S rRNA RT-PCR | 5S_Fw |
| mvRNA and 5S rRNA RT-PCR | 5S100_Rev_ATTO647N |
| TSO-based RT-PCR/qPCR | Tuni-13 LNA3 |
| TSO-based RT-PCR/qPCR | Oligo d(T)20 |
| TSO-based RT-PCR | TSO |
| TSO-based RT-PCR | TSO Fw |
| TSO-based RT-PCR | PB2 Rev |
| TSO-based RT-PCR | PB1 Rev |
| TSO-based RT-PCR | PA Rev |
| TSO-based RT-PCR | HA Rev |
| TSO-based RT-PCR | NP Rev |
| TSO-based RT-PCR | NA Rev |
| TSO-based RT-PCR | M Rev |
| TSO-based RT-PCR | NS Rev |
| TSO-based RT-PCR | VN04 PB2 TSO rev |
| TSO-based RT-PCR | VN04 HA TSO rev |
| TSO-based RT-PCR | VN04 NP TSO rev |
| TSO-based RT-PCR | VN04 NA TSO rev |
| TSO-based RT-PCR | NS H5N1 Rev |
| qPCR | 18S rRNA Fw |
| qPCR | 18s rRNA Rev |
| qPCR | NA qPCR Fw |

| | |
|---|---|
| qPCR | NA qPCR Rev |
| qPCR | PB1 qPCR Fw |
| qPCR | PB1 qPCR Rev |
| *In vitro* transcription | NA T7 ccRNA/mRNA Fw |
| *In vitro* transcription | NA T7 vRNA/svRNA Fw |
| *In vitro* transcription | NA T7 cRNA Fw |
| *In vitro* transcription | NA cRNA/ccRNA rev (NA 5′) |
| *In vitro* transcription | NA mRNA Rev |
| *In vitro* transcription | NA ccRNA-3′ Rev |
| *In vitro* transcription | NA vRNA Rev |
| *In vitro* transcription | NA svRNA Rev |
| ONT sequencing/*In vitro* transcription | NS spike T7 PCR Fw |
| ONT sequencing/*In vitro* transcription | NS spike T7 PCR Rev |
| ONT sequencing/*In vitro* transcription | NS spike gBlock DNA template for IVT |
| ONT sequencing | NS seq Fw |
| ONT sequencing | NS seq Rev |
| HEK293-luc cell line validation | HSV-1 60-mer DNA Fw |
| HEK293-luc cell line validation | HSV-1 60-mer DNA Rev |
| DVG RT-PCR/NGS/qPCR | Tuni-12 |
| DVG RT-PCR/NGS | Tuni-13 |

| Target | Step |
|---|---|
| | PCR |
| | PCR |
| | PCR |
| | PCR |
| | PCR |
| | PCR |
| | PCR |
| | PCR |
| | PCR |
| | PCR |
| | PCR |
| | PCR |
| | PCR |
| | PCR |
| | PCR |
| | PCR |
| | PCR |
| | PCR |
| 5S rRNA | RT |
| NP vRNA | RT |
| NP c/mRNA internal | RT |
| NA vRNA internal | RT |
| NA c/mRNA internal | RT |
| PB1 vRNA internal | RT |
| PB1 c/mRNA internal | RT |
| HA vRNA internal | RT |
| HA c/mRNA internal | RT |
| M vRNA internal | RT |
| M c/mRNA internal | RT |

| | |
|---|---|
| NA cRNA terminal | RT |
| NA vRNA terminal | RT |
| NA c/mRNA internal (for short templates) | RT |
| NP cRNA terminal | RT |
| NP vRNA terminal | RT |
| NP c/mRNA internal (for short templates) | RT |
| mvRNA | RT |
| mvRNA | RT |
| mvRNA | 2nd strand |
| mvRNA | PCR |
| mvRNA | PCR |
| 5S rRNA | RT |
| 5S rRNA | 2nd strand and PCR |
| 5S rRNA | PCR |
| cRNA 3' | RT |
| mRNA | RT |
| N/A | RT |
| TSO | PCR |
| PB2 A/WSN/33 | PCR |
| PB1 A/WSN/33, A/Vietnam/1203/04 | PCR |
| PA A/WSN/33, A/Vietnam/1203/04 | PCR |
| HA A/WSN/33 | PCR |
| NP A/WSN/33 | PCR |
| NA A/WSN/33 | PCR |
| M A/WSN/33, A/Vietnam/1203/04 | PCR |
| NS A/WSN/33 | PCR |
| PB2 A/Vietnam/1203/04 | PCR |
| HA A/Vietnam/1203/04 | PCR |
| NP A/Vietnam/1203/04 | PCR |
| NA A/Vietnam/1203/04 | PCR |
| NS A/Vietnam/1203/04 | PCR |
| 18S rRNA | PCR |
| 18S rRNA | RT and qPCR |
| NA | qPCR |

| | |
|---|---|
| NA | RT and qPCR |
| PB1 | qPCR |
| PB1 | qPCR |
| ccRNA/mRNA/ccRNA-3′ | PCR |
| vRNA/svRNA | PCR/annealing |
| cRNA | PCR |
| cRNA/ccRNA | PCR |
| mRNA | PCR |
| ccRNA-3′ | PCR |
| vRNA | PCR |
| svRNA | annealing |
| NS spike | PCR |
| NS spike | PCR |
| NS spike | PCR |
| NS and NS spike | PCR |
| NS and NS spike | RT and PCR |
| | annealing |
| | annealing |
| vRNA 3' | RT/PCR |
| cRNA 3' | RT/PCR |

| Sequence (5′- 3′) |
| --- |
| CTCCTGGATCCCCAAAGCAAATCGATCCATCTTG |
| CAAGATGGATCGATTTGCTTTGGGGATCCAGGAG |
| GGATCCCCAAAAGAGCTCGATCCATCTTGAATACAAGC |
| GCTTGTATTCAAGATGGATCGAGCTCTTTTGGGGATCC |
| GGATCCCCAAAAGAAATCGAGCTATCTTGAATACAAGCC |
| GGCTTGTATTCAAGATAGCTCGATTTCTTTTGGGGATCC |
| GGATCCCCAAAAGAGCTCGATCCATCTTGAATAC |
| GTATTCAAGATGGATCGAGCTCTTTTGGGGATCC |
| CATCTTGAATGCAAGCCAAAG |
| GATCGATTTCTTTTGGGG |
| CGATCCATCTTGAATACAGCTCAAAGAGGAATACTTG |
| CAAGTATTCCTCTTTGAGCTGTATTCAAGATGGATCG |
| CAGCCTAATCAGACCAAAGGAGAATCCAGCACACAAG |
| CTTGTGTGCTGGATTCTCCTTTGGTCTGATTAGGCTG |
| CTACAGGTTGTGGAGCAACCTTTGTTTCAC |
| GTGAAACAAAGGTTGCTCCACAACCTGTAG |
| AAGTTTTGAAGCAAGAGCAGCGATATTCTGTGCCCGACAGAACTGC |
| GAAAACTGCGCTGGCTTGGGATGTGGTCTACTCACAAAGCATTCC |
| TCCCAGGCGGTCTCCCATCC |
| ATTTCTTCGGAGACAATGCAG |
| TAAGATCGTTTGGTGCCTTTG |
| TGGACTAGTGGGAGCAT |
| TCCAGTATGGTTTTGATTTCC |
| TGATTTCGAATCTGGAAGGA |
| TCCATGGTGTATCCTGTTCC |
| TACTCAACTGTCGCCAGTTC |
| GTCACTGCCACATTCTTCTC |
| GAAAAGAGGGCCTTCTACGG |
| AGCCATTCCATGAGAACCTC |

+ for LNA bases
H for Adenine, Cytosine or Thymine
N for Adenine, Guanine, Cytosine or Thymi[illegible]
r for ribonucleotides

| |
|---|
| AGTAGAAACAAGGAGTTTTTTG |
| AGCGAAAGCAGGAGTTTAAATG |
| CTACTTGTCAATGGTGAACG |
| AGTAGAAACAAGGGTATTTTTC |
| AGCAAAAGCAGGGTAGACTAGT |
| GTCGTACTCCTCTGCATTG |
| GTTCAGACGTGTGCTCTTCCGATCTAGC+A+AAAGCAGG |
| GTTCAGACGTGTGCTCTTCCGATCTAGCG+AAAGCAGG |
| CACGACGCTCTTCCGATCTHNNNNNNNAGTAGAA+A+CAAGG |
| GACGTGTGCTCTTCCGATCT |
| CACGACGCTCTTCCGATCT |
| TCCCAGGCGGTCTCCCATCC |
| GTCTACGGCCATACCACC |
| TCCCAGGCGGTCTCCCATCC |
| ACGCGTGATCAGTAGAAA+CA+AG+G |
| TTTTTTTTTTTTTTTTTTTT |
| GCTAATCATTGCAAGCAGTGGTATCAACGCAGAGTACATrGrGrG |
| CATTGCAAGCAGTGGTATCAAC |
| CTGCGACATTAGATTCCTTAGTTC |
| GTAAAGTCGGATTGACATCCATTC |
| GGATTGAAGCATTGTCGCAC |
| CAGGACTAGTACAAAAGCCTTC |
| GATTTCGATGTCACTCTGTGAG |
| GATCCAATGGTTATTATTTTCTGGTTTG |
| GACCTCGGTTAGAAGACTCATC |
| GCTTGACACAGTGTTTGGATC |
| CTGTGACATTAGATCTCGTAATTC |
| CACTTTTAACAAGACTGACTATTGC |
| CGTTTGGTGCCTTGAGACG |
| GATCCGATGGTTATTATCTTCTG |
| GCGGACATGCCAAAGAAAG |
| ACCCGTTGAACCCCATTGGTGA |
| GCCTCACTAAACCATCCAATCGG |
| GTTTGAATCGGTTGCTTGG |

CTGCTCCATCATCTGGACC
CTGTATTAGGCGTCTCCATC
CATCCCACCAGTAAGTAGTC
TAATACGACTCACTATAGGGAATACTCAAGGCAAAAGCAGGAGTTTAAATG
TAATACGACTCACTATTAGTAGAAACAAGGAGTTTTTTGAAC
TAATACGACTCACTATTAGCAAAAGCAGGAGTTTAAATG
AGTAGAAACAAGGAGTTTTTTG
TTTTTTTTTTTTTTTTTTTTTTTTTTTTTTGAACAAACTACTTGTCAATGGTG
AAACTACTTGTCAATGGTGAACG
AGCAAAAGCAGGAGTTTAAATG
GTTCAAAAAACTCCTTGTTTCTACTAATAGTGAGTCGTATTA
TAATACGACTCACTATAGGGAATACTC
TCCTTGTTAGAGGGTGTTTTTTAT
TAATACGACTCACTATAGGGAATACTCAAGGCAAAAGCAGGGTGACAAAGACATAATGGATCCAAACACTGTGTCAAGCTT
GACTTCTGATCTCGGCGAAG
ACTCTCGGTCTGGACATC
TAAGACACGATGCGATAAAATCTGTTTGTAAAATTTATTAAGGGTACAAATTGCCCTAGC
GCTAGGGCAATTTGTACCCTTAATAAATTTTACAAACAGATTTTATCGCATCGTGTCTTA
ACGCGTGATCAGCAAAAGCAGG
ACGCGTGATCAGTAGAAACAAGG

ne

TCAGGTAGATTGCTTTCTTTGGCATGTCCGCAAAAGAGTTGCAGACCAAGAACTAGGTGATGCCCCATTATCCAC

CGTGCTGGAAAGCAAATAGTGGAGCGGATTCTGAAGGAAGAATCTGATGAGGCACTCAAAATGACCATGGCCTC

TGTACCTGCATCGCGCTACCTAACTGACATGACTCTTGAGGAAATGTCAAGGCACTGGTTCATGCTCATGCCCAA

AGCAGAAAGTGGCAGGCCCTCTTTGTATCAGAATGGACCAGGCGATCATGGATAAGAACATCATACTGAAAGCGA

AACTTCAGTGTGATTTTTGACCGGCTGGAGACTCTAATATTACTAAGGGCCTTCACCGAAGAGGGGACAATTGTT[illegible]

GGCGAAATTTCACCACTGCCCTCTCTTCCAGGACATACTGATGAGGATGTCAAAAATGCAGTTGGGGTCCTCATC

TGAGAATGGGAGACCTCCACTCACTCCAAAACAGAAACGGAAAATGGCGGGAACAATTAGGTCAGAAGTTTGAA[illegible]

GAAATAAGATGGTTGATTGAAGAAGTGAGACACAGACTGAAGATAACAGAGAATAGTTTTGAGCAAATAACATTTA

ATGCAAGCCTTACAACTATTGCTTGAAGTGGAGCAAGAGATAAGAACTTTCTCGTTTCAGCTTATTTAATAATAAAA

AACACCCTCTAACAAGGA

**Supplementary Tabl**

| Template name |
| --- |
| NA76 |
| NA124 |
| NA196 |
| NA244 |
| NP76 |
| NP125 |
| NP197 |
| NP246 |

**e 8. Sequences of internally truncated vRNA templates used in RNP assays**

| vRNA-sense sequence (5′- 3′) |
| --- |
| AGUAGAAACAAGGAGUUUUUUGAACAAACUACUUGUCAAUUUCUGGUUUGGAUUCAUUUA |
| AGUAGAAACAAGGAGUUUUUUGAACAAACUACUUGUCAAUGGUGAACGGGAGCUCAGCA[illegible] |
| AGUAGAAACAAGGAGUUUUUUGAACAAACUACUUGUCAAUGGUGAACGGGAGCUCAGCA[illegible] |
| AGUAGAAACAAGGAGUUUUUUGAACAAACUACUUGUCAAUGGUGAACGGGAGCUCAGCA[illegible] |
| AGUAGAAACAAGGGUAUUUUUCUUUACUAGUGACUUCGAUGUCACUCUGUGAGUACUAG |
| AGUAGAAACAAGGGUAUUUUUCUUUAAUUGUCGUACUCCUCUGCAUUGUCACUAGUUCG |
| AGUAGAAACAAGGGUAUUUUUCUUUAAUUGUCGUACUCCUCUGCAUUGUCUCCGAAGAAA |
| AGUAGAAACAAGGGUAUUUUUCUUUAAUUGUCGUACUCCUCUGCAUUGUCUCCGAAGAAA |

AAACUCCUGCUUUUGCU

CCGUACAGAUCGAUCCAAUGGUUAUUAUUUUCUGGUUUGGAUUCAUUUAAACUCC

CCGUCUGGCCAAGACCAAUCUACAGUAUCACCAUUCACACUAUUUGCAAUAUUAG

CCGUCUGGCCAAGACCAAUCUACAGUAUCACCAUUCACACCACAAAAAGAAAUGA

UCUACCCUGCUUUUGCU

UUUGGUGCCUUUGGUCGCCAUGAUUUCGAUGUCACUCUGUGAGUACUAGUCUAC

AUAAGAUCCUUCAUUACUCAUGUCAAAGGAGGGCACGAUCGGGCUCGUUGCACU

AUAAGAUCCUUCAUUACUCAUGUCAAAGGAGGGCACGAUCGGGCUCGUUGCCUU

CUGCUUUUGCU
GGCUAAUUAUUCCGACUACCAUACAGAUCGAUCCAAUGGUUAUUAUUUUCUGGUU
UGCUCCCACAAUCCAUAUUGAGAUUAUAUUUCCUAUUUGCAAUAUUAGGCUAAUU

CCCUGCUUUUGCU
AGUCUGUUCGUAAGAUCGUUUGGUGCCUUUGGUCGCCAUGAUUUCGAUGUCACU
UUCGUCCGAGAGCUCGAAGACUCCCCGCCCCUGGAAAGACACUAGUCUCCAUCU

UGGAUUCAUUUAAACUCCUGCUUUUGCU
JAUUCCGACUACCAUACAGAUCGAUCCAAUGGUUAUUAUUUUCUGGUUUGGAUUC

JCUGUGAGUACUAGUCUACCCUGCUUUUGCU
IGUUCGUAAGAUCGUUUGGUGCCUUUGGUCGCCAUGAUUUCGAUGUCACUCUGU

CAUUUAAACUCCUGCUUUUGCU

GAGUACUAGUCUACCCUGCUUUUGCU

**Supplementary Table 9. RNAs used in this study**

| **Experiment** |
|---|
| transcription assay |
| transcription assay |
| transcription assay |
| cap cleavage assay |
| cap cleavage assay |
| cap cleavage assay |
| *In vitro* transcription/RNA transfection/TSO-based RT PCR optimisation |
| *In vitro* transcription/RNA transfection/TSO-based RT PCR optimisation |
| *In vitro* transcription/RNA transfection/TSO-based RT PCR optimisation |
| *In vitro* transcription/RNA transfection/TSO-based RT PCR optimisation |
| *In vitro* transcription/RNA transfection/TSO-based RT PCR optimisation |
| *In vitro* transcription/RNA transfection/TSO-based RT PCR optimisation |
| ONT sequencing |

| Name |
| --- |
| AG11_primer for capping |
| NA71 |
| NA71-U |
| 20-mer RNA with 5' diphosphate |
| vRNA_5p |
| vRNA_3p |
| NA196 vRNA |
| NA196 cRNA |
| NA196 mRNA |
| NA196 ccRNA |
| NA196 ccRNA-3′ |
| NA svRNA |
| NS spike |

| **sequence (5′- 3′)** |
|---|
| ppGAAUACUCAAG |
| AGUAGAAACAAGGAGUUUUUUGAACAAACUACUUGUCAAUGGUUGGAUUCAUUUAAACU[illegible] |
| AGUAGAAACAAGGAGAUGUGUGAACAAACUACUUGUCAAUGGUUGGAUUCAUUUAAACU[illegible] |
| pp AAUCUAUAAUAGCAUUAUCC |
| AGUAGAAACAAGGCC |
| GGCCUGCUUUUGCU |
| AGUAGAAACAAGGAGUUUUUUGAACAAACUACUUGUCAAUGGUGAACGGGAGCUCAGCA[illegible] |
| AGCAAAAGCAGGAGUUUAAAUGAAUCCAAACCAGAAAAUAAUAACCAUUGGAUCGAUCUG |
| GGGAAUACUCAAGGCAAAAGCAGGAGUUUAAAUGAAUCCAAACCAGAAAAUAAUAACCAU[illegible] |
| GGGAAUACUCAAGGCAAAAGCAGGAGUUUAAAUGAAUCCAAACCAGAAAAUAAUAACCAU[illegible] |
| GGGAAUACUCAAGGCAAAAGCAGGAGUUUAAAUGAAUCCAAACCAGAAAAUAAUAACCAU[illegible] |
| AGUAGAAACAAGGAGUUUUUUGAAC |
| GGGAAUACUCAAGGCAAAAGCAGGGUGACAAAGACAUAAUGGAUCCAAACACUGUGUCAA[illegible] |

CCUGCUUUUGCU
CCUGCUUUUGCU

CCGUCUGGCCAAGACCAAUCUACAGUAUCACCAUUCACACUAUUUGCAAUAUUAG
UAUGGUAGUCGGAAUAAUUAGCCUAAUAUUGCAAAUAGUGUGAAUGGUGAUACU
UGGAUCGAUCUGUAUGGUAGUCGGAAUAAUUAGCCUAAUAUUGCAAAUAGUGUG
UGGAUCGAUCUGUAUGGUAGUCGGAAUAAUUAGCCUAAUAUUGCAAAUAGUGUG
UGGAUCGAUCUGUAUGGUAGUCGGAAUAAUUAGCCUAAUAUUGCAAAUAGUGUG

AGCUUUCAGGUAGAUUGCUUUCUUUGGCAUGUCCGCAAAAGAGUUGCAGACCAA

GCUAAUUAUUCCGACUACCAUACAGAUCGAUCCAAUGGUUAUUAUUUUCUGGUU
GUAGAUUGGUCUUGGCCAGACGGUGCUGAGCUCCCGUUCACCAUUGACAAGUA
AAUGGUGAUACUGUAGAUUGGUCUUGGCCAGACGGUGCUGAGCUCCCGUUCAC
AAUGGUGAUACUGUAGAUUGGUCUUGGCCAGACGGUGCUGAGCUCCCGUUCAC
AAUGGUGAUACUGUAGAUUGGUCUUGGCCAGACGGUGCUGAGCUCCCGUUCAC

GAACUAGGUGAUGCCCCAUUAUCCACCACCCUUCGCCGAGAUCAGAAGUCCCUA

UGGAUUCAUUUAAACUCCUGCUUUUGCU
GUUUGUUCAAAAAACUCCUUGUUUCUACU
CAUUGACAAGUAGUUUGUUCAAAAAAAAAAAAAAAAAAAAAAAAAAAAAA
CAUUGACAAGUAGUUUGUUCAAAAAACUCCUUGUUUCUACU
CAUUGACAAGUAGUUU

AGAGGAAGAGGCAGCACUCUCGGUCUGGACAUCUGUCAGAAGCCCCGUGCUGGA

AAAGCAAAUAGUGGAGCGGAUUCUGAAGGAAGAAUCUGAUGAGGCACUCAAAAUG

UUGUAUCAGAAUGGACCAGGCGAUCAUGGAUAAGAACAUCAUACUGAAAGCGAAC

CGAAGAGGGGACAAUUGUUGGCGAAAUUUCACCACUGCCCUCUCUUCCAGGACAU

JACUGAUGAGGAUGUCAAAAAUGCAGUUGGGGUCCUCAUCGGAGGACUUGAAUG

CAGUAAUGAGAAUGGGAGACCUCCACUCACUCCAAAACAGAAACGGAAAAUGGC[illegible]

GGGAACAAUUAGGUCAGAAGUUUGAAGAAAUAAGAUGGUUGAUUGAAGAAGUGAG

GACACAGACUGAAGAUAACAGAGAAUAGUUUUGAGCAAAUAACAUUUAUGCAAGCC

JAUUUAAUAAUAAAAAACACCCUCUAACAAGGA
