## Supplemental Figures for "Aberrant influenza virus transcription termination generates capped cRNAs that activate RIG-I"

### Supplementary Figures 1-7

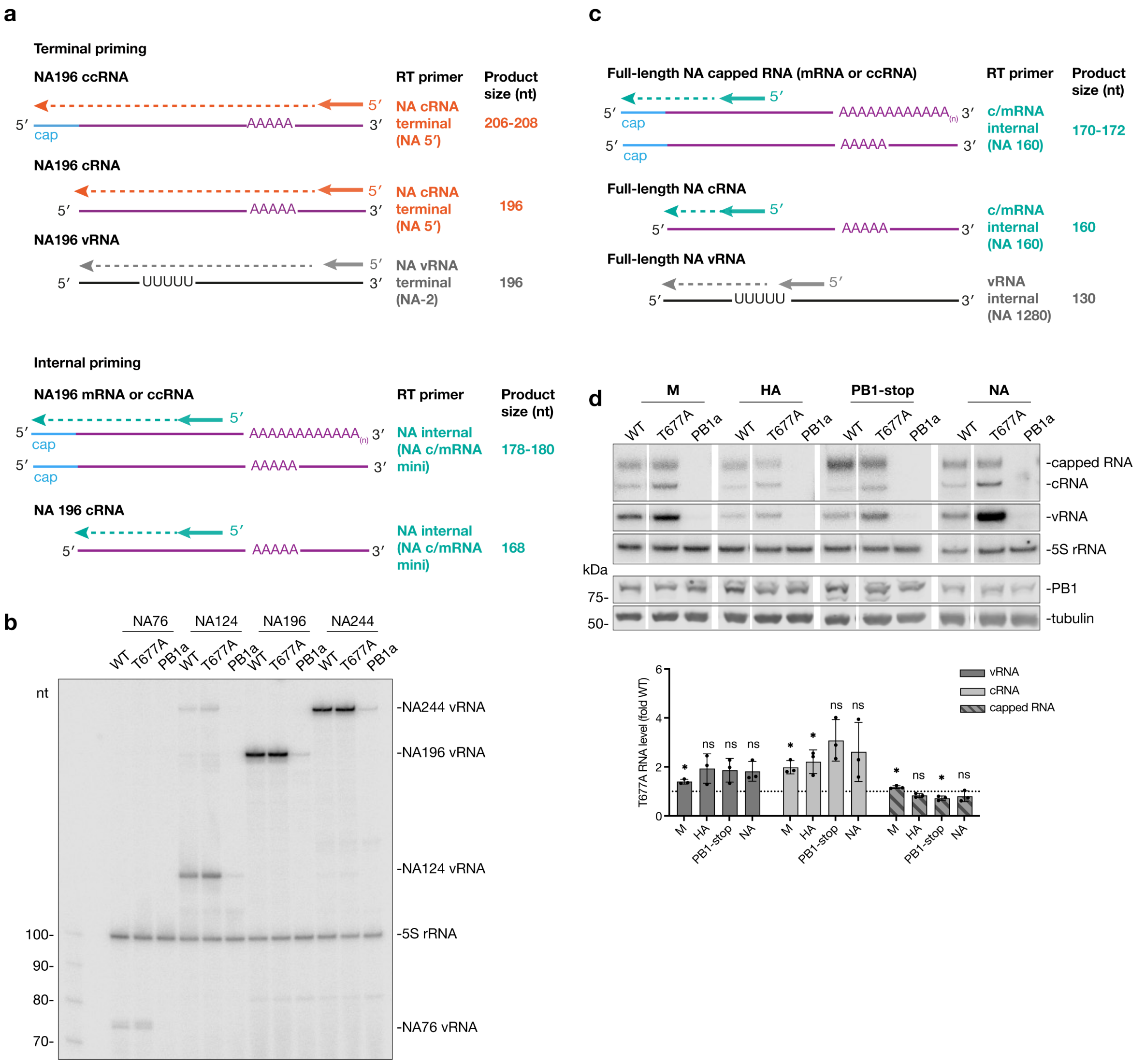


**Supplementary Fig. 1. RNP reconstitution assay with full-length and short templates. a**, Schematic of primer extension assay using terminal or internal primers for short segment 6-based templates using the NA196 template as an example. Primer binding sites and sizes of expected products are indicated. **b**, Primer extension analysis of vRNA (using NA-2 primer) in RNP reconstitution assay with short segment 6-based templates (quantification shown in Fig. 1e). **c**, Schematic of the primer extension assay for the full-length segment 6 (NA) using internal c/mRNA and vRNA primers. **d**, Primer extension analysis for full-length M, HA, PB1 with a start codon mutation (PB1-stop), and NA templates and western blot analysis of protein expression. Inactive polymerase mutant (PB1a) was used as a negative control. Primers used were: M vRNA and M c/mRNA; HA vRNA and HA c/mRNA; PB1 vRNA and PB1 c/mRNA; NA 1280 and NA 160. Quantification of vRNA, cRNA and all capped RNA levels produced by the T677A mutant versus wild-type polymerase is shown at the bottom. All gel images are representative of three independent replicates. Data in panel **d** is presented as mean ± SD from three independent experiments. Statistical significance was determined using a one-sample t-test, where each condition was compared to a theoretical mean of 1 (ns=non-significant, **P*<0.05).

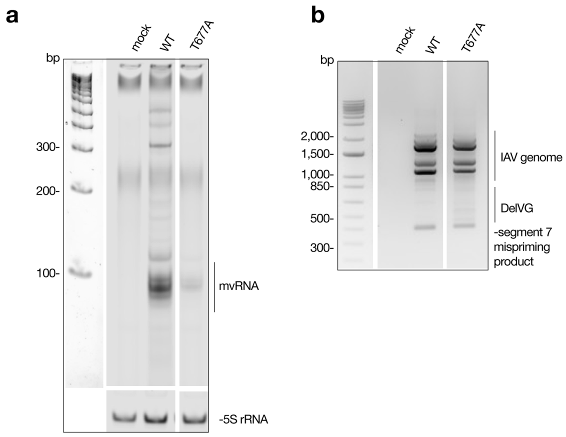


**Supplementary Fig. 2. Generation of non-canonical replication products by wild-type and T677A viruses.** RT-PCR analysis of mvRNA synthesis was performed using RNA extracted from HEK293-luc cells infected with the wild-type or T677A viruses at an MOI of 0.01 for 48 hours. Total RNA was size-fractionated, and the small fraction (<200 nt) was used for RT-PCR with universal primers targeting the conserved genome segment termini. 5S rRNA was used as a cellular RNA loading control. **b**, DelVG production was assessed in the same infection samples as in panel **a**, using total RNA and RT-PCR with universal primers. All gel images are representative of three independent replicates.

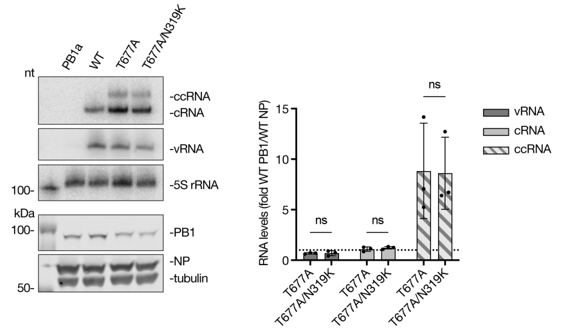


**Supplementary Fig. 3. RNP reconstitution assay with PB1 T677A/NP319K double mutant.** RNP assay was performed using wild-type PB1 and NP, T677A PB1 and wild-type NP or a PB1 T677A/NP N319K double mutant, with the NA196 template. Primer extension analysis was done using primers targeting the 3' termini of cRNA/ccRNA (NA 5′) and vRNA (NA-2) molecules. Protein expression was assessed with a western blot. Gel image is representative of three replicates. Data are shown as mean ± SD from three independent experiments. Statistical significance was determined using two-way ANOVA with with Šídák's multiple comparisons test (ns=non-significant).

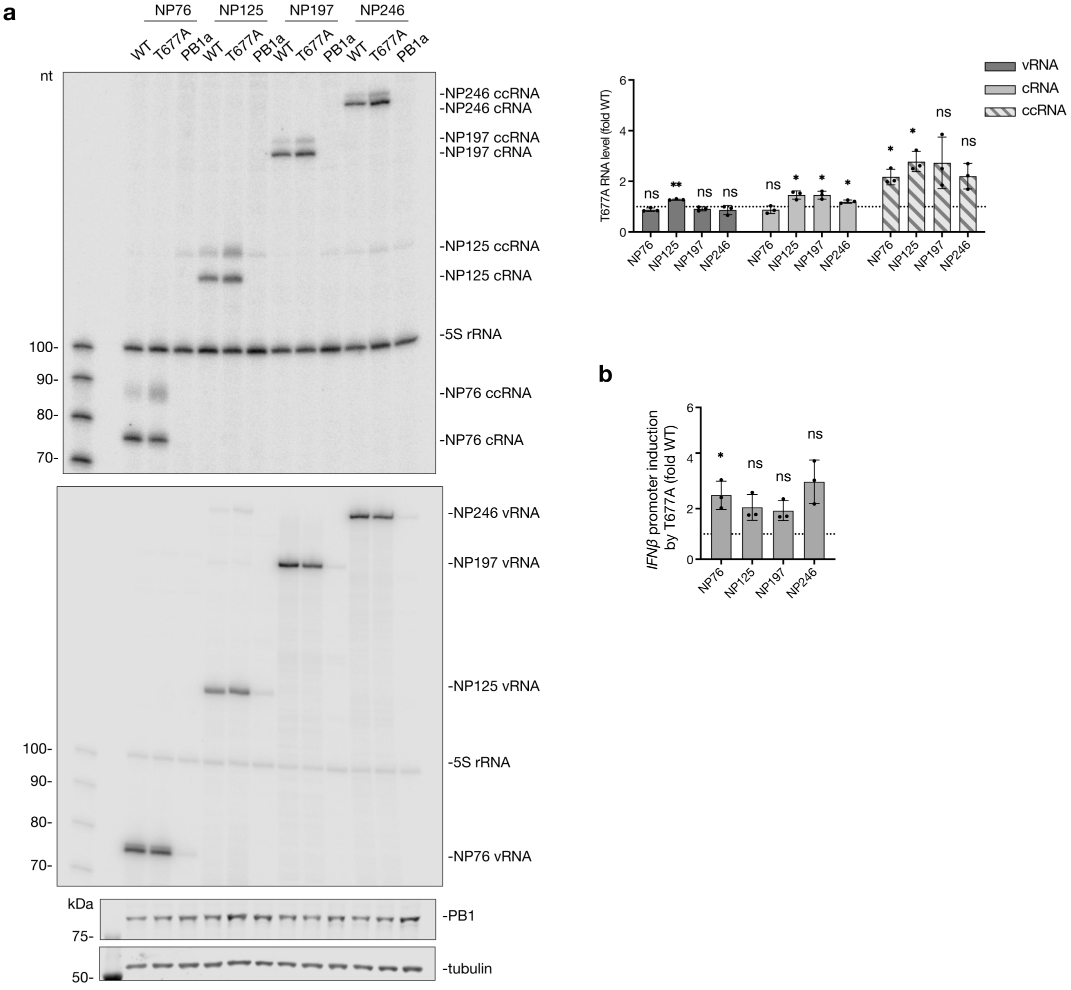


**Supplementary Fig. 4. RNP reconstitution assays with short segment 5-based templates. a**, Primer extension analysis using the terminal cRNA (NP 5′) and vRNA (GC-) primers. Representative primer extension gel images (top two images) and western blot analysis of protein expression (bottom two images) from three independent experiments are shown. The quantification of the vRNA, cRNA and ccRNA levels produced by the T677A relative to the wild-type polymerase are shown in the top right graph. **b**, Quantification of *IFN-β* promoter activity in the RNP assay with segment-5 based templates. Data in panels a and b are shown as mean ± SD from three independent experiments. Statistical significance was determined using a one-sample t-test where each condition was compared to a theoretical mean of 1 (ns=non-significant, *$P$<0.05, **$P$<0.01).

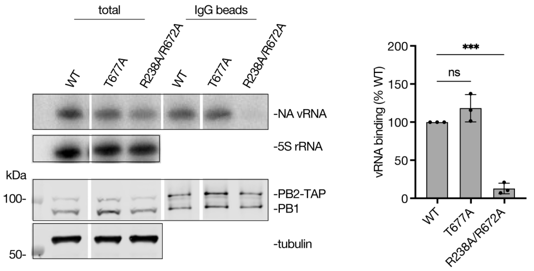


**Supplementary Fig. 5. Polymerase-vRNA binding assay.** Polymerases containing a PB2-TAP tag and either wild-type PB1, T677A PB1 or the R238A/R672A PB1 (incapable of promoter binding and used as a negative control), were co-expressed with a full-length segment 6 (NA) vRNA template in HEK293T cells. Polymerases were captured on IgG beads and levels of co-precipitating NA vRNA were assessed by primer extension analysis with NA 1280 primer (right). Gel image is representative of three independent experiments and data are presented as mean ± SD. Statistical significance was determined using one-way ANOVA with Dunnett's multiple comparisons test (ns=non-significant, ****P*<0.001).

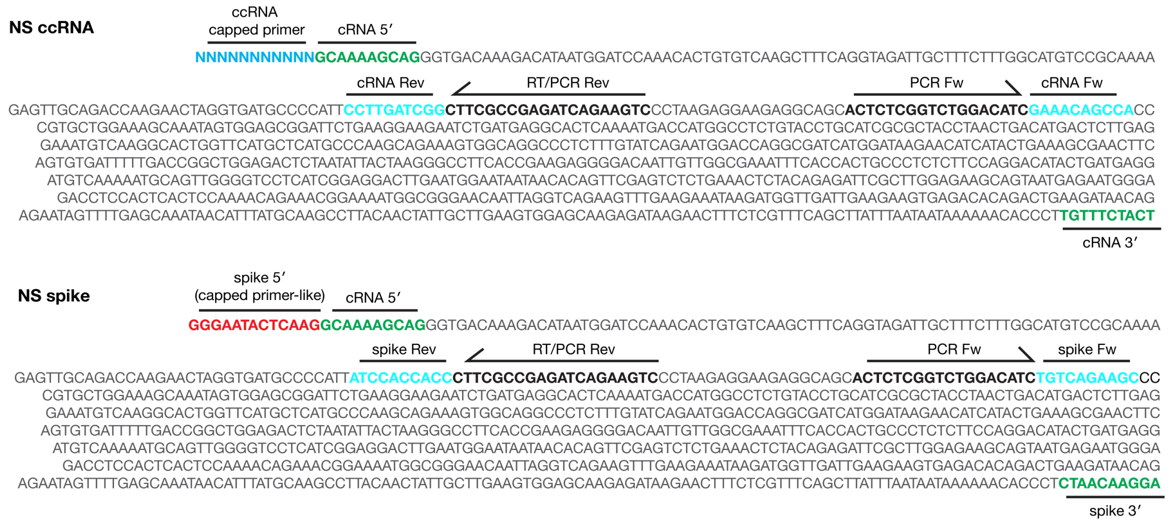


**Supplementary Fig. 6. Schematic of NS ccRNA and NS spike.** Capped ccRNA primer is shown in blue; spike 5′ capped primer-like sequences in red; cRNA 5′ and 3′ termini and the spike 3′ barcode in green. RT-PCR primer binding sites are shown in bold and primer-adjacent spike barcodes or cRNA sequences are shown in cyan.

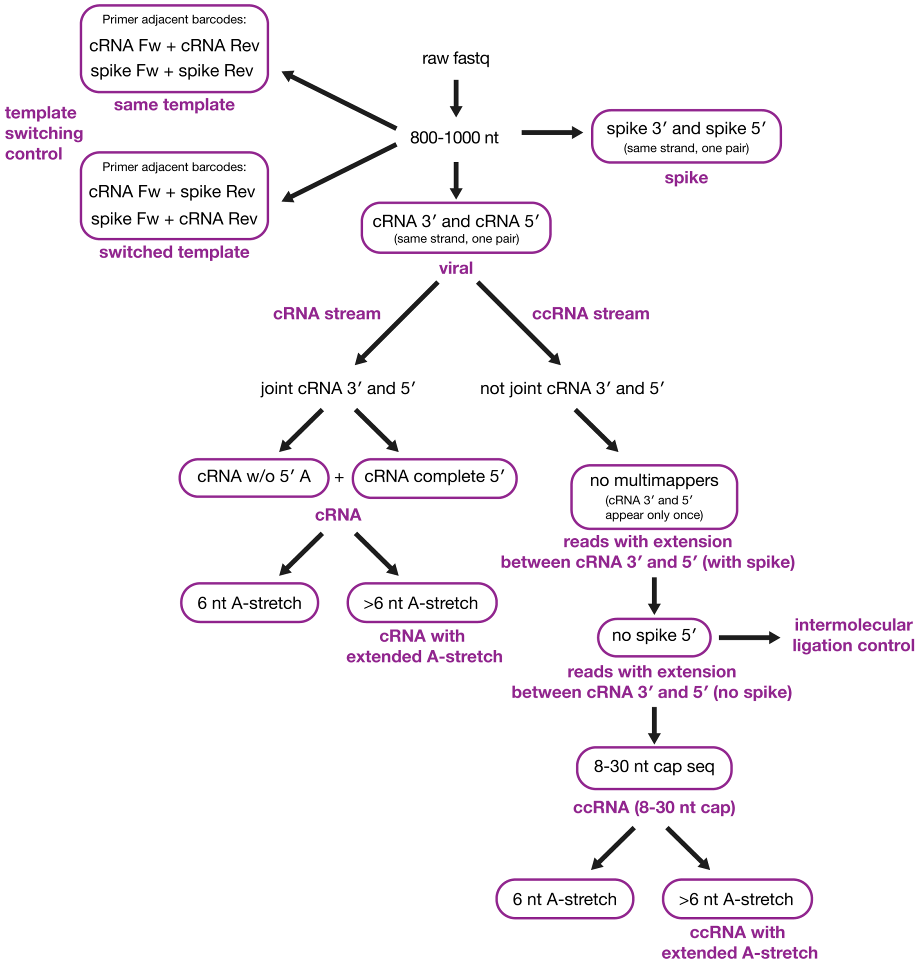


**Supplementary Fig. 7. Schematic of ONT sequencing data analysis.** Description of the schematic is provided in the Supplementary Note.

### Supplementary Note

*Description of ONT sequencing analysis*

To minimize off-target or degraded fragments, raw Oxford Nanopore sequencing reads were size-selected; reads shorter than 800 nt or longer than 1000 nt were excluded from downstream analysis. The remaining reads (800 – 1000 nt) were screened for the presence of viral cRNA 3' terminal (TGTTTCTACT) and viral cRNA 5' terminal (GCAAAAGCAG) motifs (Supplementary Fig. 5). Only reads possessing both motifs in the same sense orientation were retained. Next, cRNAs and ccRNAs were routed into separate analysis streams based on the presence of a specific cRNA junction sequence. Reads containing joint cRNA 3' and 5' ends with or without the first adenosine at cRNA 5' (either TGTTTCTACTAGCAAAAGCAG or TGTTTCTACTGCAAAAGCAG) were routed into cRNA stream, while reads lacking these exact junction sequences were channeled into the ccRNA stream. To isolate and extract host-derived capped primer sequences from the processed ccRNA pools, reads were reevaluated for the viral cRNA 3' and 5' termini under the additional constraint that each motif must appear in the same sense orientation and must only appear once. Reads containing multiple copies of either motif were discarded to eliminate ambiguity in the extracted capped primer region. The number of capped primer sequences containing spike 5' sequence (GGGAATACTCAAG) were quantified to estimate intermolecular ligation rate and subsequently removed from the ccRNA pool. Reads with 8-30 nt extensions between 3' and 5' cRNA termini were considered legitimate ccRNAs.

To measure the length of the A-stretch extension, cRNAs and ccRNAs with a validated capped primer length (8-30 nt) were screened for the presence of the motifs that flank the A-stretch (ATTTAATAAT and CACCCTTGTT). The intervening sequence between these two motifs was isolated to determine the exact length of the A-stretch. As with previous filtering steps, reads containing multiple copies of either flanking motif were discarded to prevent ambiguity. Any sequences not consisting solely of adenines were also discounted.

To determine the spike to viral reads ratio all size-selected reads were screened for sequences possessing both spike 3' barcode (CTAACAAGGA) and spike 5' terminus (GGGAATACTCAAG). The resulting number of reads was divided by those containing viral cRNA 3' and 5' ends.

To determine the rate of template switching, we searched for reads containing primer adjacent barcodes coming from the same template (i.e. cRNA Fw (GAAACAGCCA) and cRNA Rev (CCTTGATCGG) or spike Fw (TGTCAGAAGC) and spike Rev (ATCCACCACC)) or coming from the switched templates (cRNA Fw and spike Rev or spike Fw and cRNA Rev). Then we divided the total number of switched template reads by the number of sample template reads.
