## Extended Figures for "Aberrant influenza virus transcription termination generates capped cRNAs that activate RIG-I"

### Extended Data

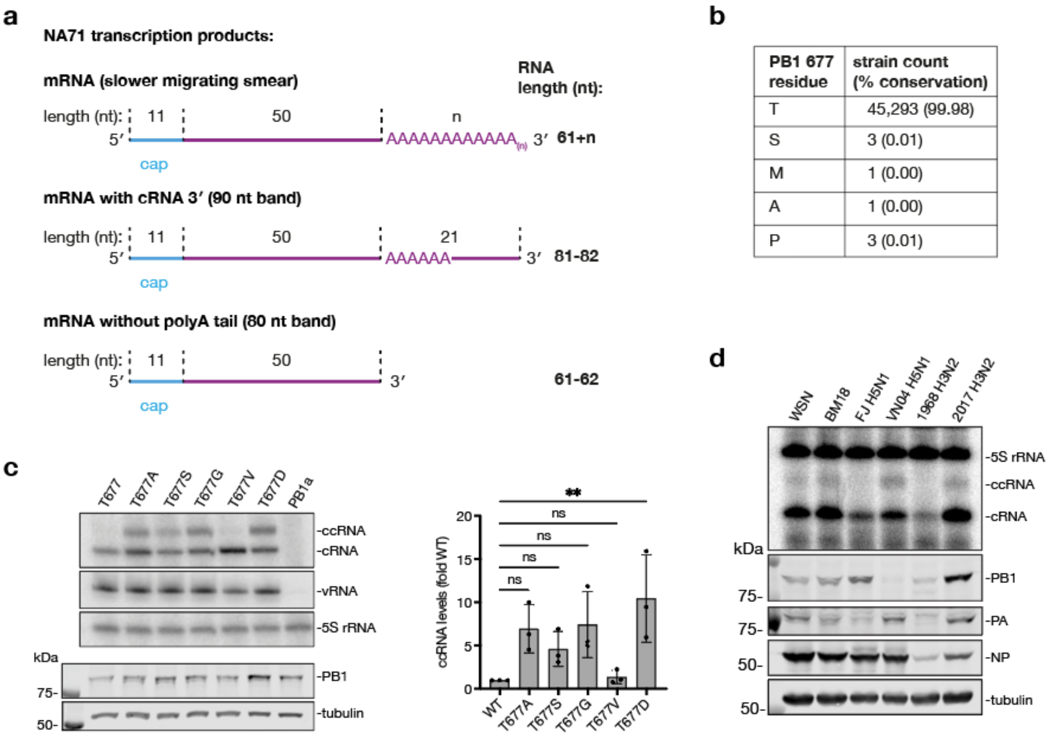


| PB1 677 residue | strain count (% conservation) |
|---|---|
| T | 45,293 (99.98) |
| S | 3 (0.01) |
| M | 1 (0.00) |
| A | 1 (0.00) |
| P | 3 (0.01) |

**Extended Data Fig. 1| *In vitro* activity assays with purified polymerases and RNP reconstitution assays. a**, Schematic of proposed transcription products observed in Fig. 1b, c. **b**, Amino acid variation at position 677 of IAV PB1 (analysis done using www.fludb.org database). **c**, RNP reconstitution assay using the NA196 template and A/WSN/33 polymerases with different PB1 T677 amino acid substitutions. Primer extension analysis with terminal cRNA (NA 5′) and vRNA (NA-2) primers (top three images) and western blot analysis of protein expression levels are shown. The quantification of ccRNA levels is shown at the bottom. **d**, RNP assay analysis with the segment 6-derived NA76 template and the RNA polymerases of the IAV strains: A/WSN/1933 H1N1 (WSN), A/Brevig Mission/1/1918 H1N1 (BM18), A/Fuijan/1/1995 H5N1 (FJ H5N1), A/Vietnam/1203/2004 H5N1 (VN04 H5N1), A/Bilthoven/16190/1968 H3N2 (1968 H3N2), and A/Netherlands/499/2017 H3N2 (2017 H3N2). Primer extension analysis was done with the terminal cRNA primer (NA 5′) and is representative of three experiments. Viral protein expression was assessed with western blot. All gel images are representative of three independent replicates. Data in panel **c** are shown as mean ± SD from three independent experiments. Statistical significance was determined using a one-way ANOVA with Dunnett's multiple comparisons test; (ns=non-significant, **P*<0.05).

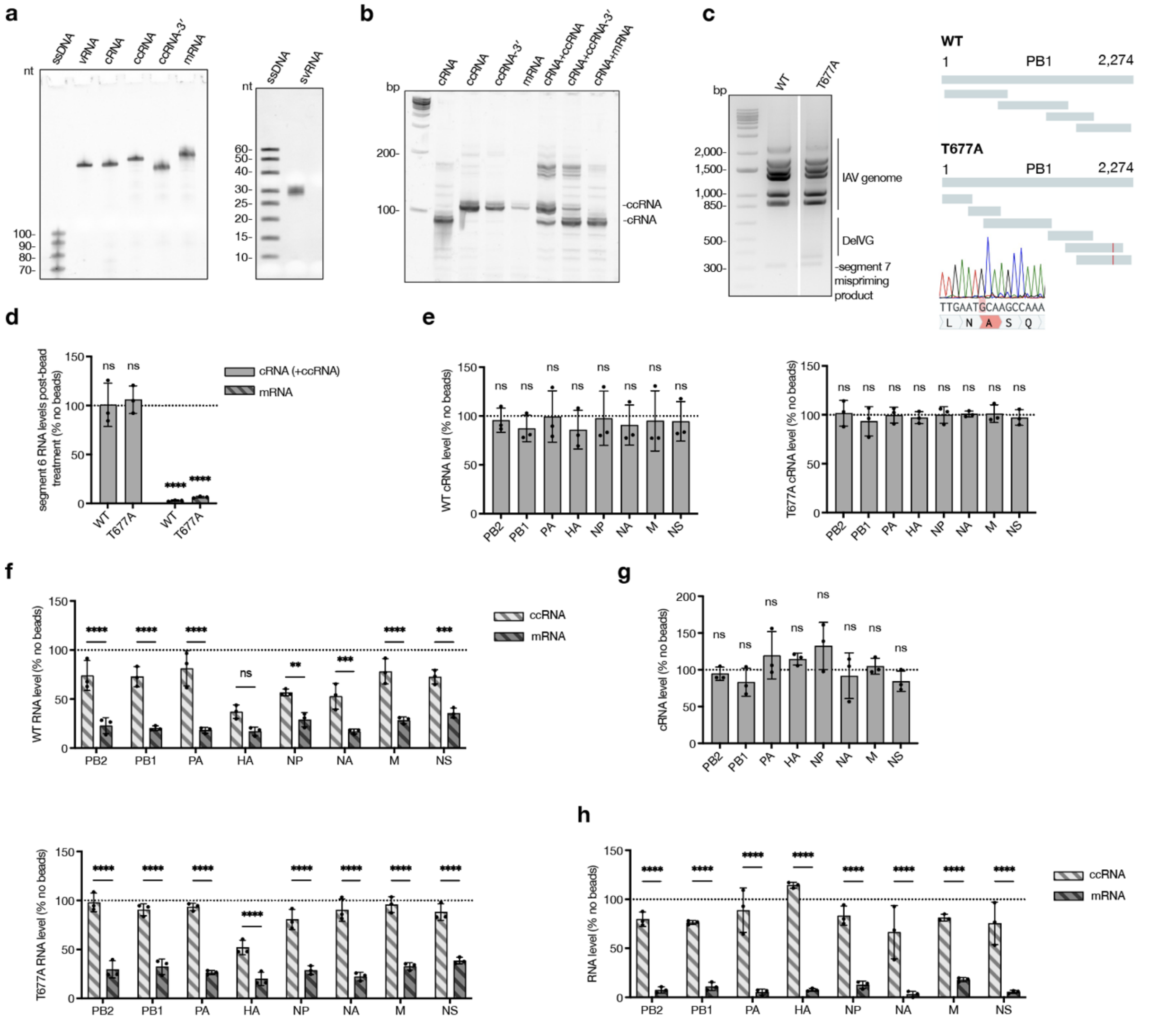


**Extended Data Fig. 2 TSO-based RT PCR optimisation and ccRNA detection in infection. a**, *In vitro* transcripts used in this study. NA196 template was used to generate vRNA, cRNA, ccRNA, ccRNA lacking 25 3'-terminal nt (ccRNA-3'), and mRNA molecules using T7 RNA polymerase. svRNA was generated using last 25 nt of segment 6 vRNA 3' terminus as a template. All transcripts were gel purified. ccRNA, ccRNA-3' and mRNA molecules were capped and methylated, and any remaining uncapped RNA enzymatically digested. The RNA quality was evaluated using denaturing PAGE with 100 ng of RNA per lane (image is shown). **b**, Products of TSO-based RT-PCR where *in vitro* transcripts (shown in a) were used as input RNA alone or in combination. Image is representative of three replicates. **c**, Analysis of wild-type and T677A virus preparations. Virus stocks were generated by reverse-genetics and plaque purified. RT-PCR was performed on viral RNA isolated from 12 individual plaques using universal Tuni-12 and Tuni-13 primers, which bind to 3' vRNA and cRNA termini, respectively. Virus isolates with the lowest DelVG content (image representative of three replicates, left) were selected and used for subsequent infection experiments. The 300 bp band observed was isolated, sequenced and identified as a mispriming product of the Tuni-12 primer to an internal sequence of segment 7. PB1 gene sequences were validated using Sanger sequencing (right). The schematic shows the areas covered by different sequence runs. The sequence trace of the T677A virus is shown at the bottom. **d**, Assessment of efficiency of mRNA depletion by oligo $d(T)_{25}$-bead treatment. Total RNA from A549 cells infected with wild-type or T677A viruses was subjected to oligo $d(T)_{25}$-bead treatment or left untreated. Segment 6 cRNA and mRNA levels were quantified by qRT-PCR, using a terminal cRNA primer (which also detects ccRNA) and oligo $d(T)_{20}$ primer, respectively. **e**, Quantification of cRNA levels following oligo $d(T)_{25}$-bead treatment of RNA extracted from wild-type and T677A virus infected A549 cells as shown in Fig. 2b. **f**, Quantification of ccRNA and mRNA levels after oligo $d(T)_{25}$-bead treatment of RNA extracted from wild-type and T677A virus infected A549 cells as shown in Fig. 2b. **g**, Quantification of cRNA levels following oligo $d(T)_{25}$-bead treatment of RNA extracted from A/Vietnam/1203/04 (H5N1) virus infected murine lungs as shown in Fig. 2d. **h**, Quantification of ccRNA and mRNA levels following oligo $d(T)_{25}$-bead treatment of RNA extracted from A/Vietnam/1203/04 (H5N1) virus infected murine lungs as shown in Fig. 2d. Gel image in panel **a** is representative of a single experiment where *in vitro* transcribed RNA used in this study was generated; images in panels **b** and **c** are representative of three independent replicates. Data in **d**-**h** are shown as mean ± SD from three independent experiments (**d**-**f**) or three infected mice (**g**, **h**). Statistical significance in **d**, **e** and **g** was determined using a one-sample t-test where each condition was compared to a theoretical mean of 1 or 100 (shown as a dotted line; grouping is for visual purposes only, no comparisons have been made between groups); statistical significance in **f** and **h** was determined using two-way ANOVA with Šídák's multiple comparisons test; (ns=non-significant, *$P<0.05$, **$P<0.01$, ***$P<0.001$, ****$P<0.0001$).

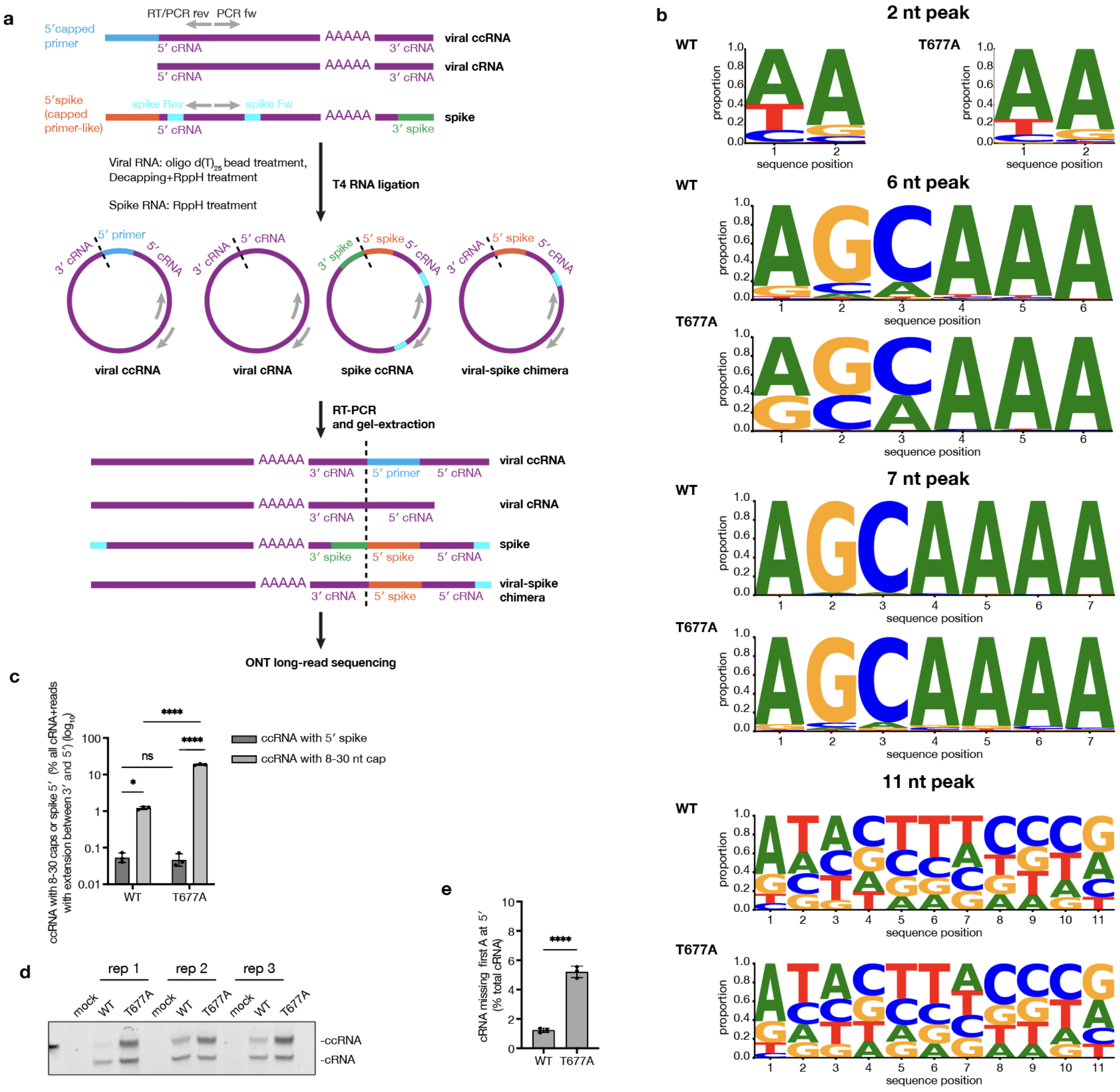


**Extended Data Fig. 3 Segment eight (NS) ccRNA detection using ONT. a,** Schematic of sample preparation for sequencing. Total cellular RNA, isolated from infected cells was treated with oligo d(T)$_{25}$ beads to remove mRNAs, then 5' m$^7$G caps and 5' pyrophosphates were removed using mRNA decapping enzyme and RNA 5' Pyrophosphohydrolase (RppH) respectively. Segment 8-based ccRNA-like spike RNA contained a specific capped primer-like sequence at the 5' terminus and a barcode at the 3' terminus. Spike RNA was treated with RppH to remove 5' pyrophosphates and added to total RNA for circularization reaction using T4 RNA Ligase 1. Positive-sense segment 8 RNA were amplified across the 3' - 5' junction using RT-PCR with segment 8-specific primers. The PCR products were gel-purified and sequenced using ONT. **b**, Position-specific nucleotide distribution for the 2-nt, 6-nt, 7nt and 11-nt ccRNA peaks for wild-type and T677A virus infection samples (data represents the sum of three replicates). **c**, Levels of ccRNAs with 8-30 nt-long capped primer extensions and those with spike 5' shown as percentage of total of cRNA and reads with extensions between cRNA 3' and 5' (before spike removal). **d**, TSO-based RT-PCR for segment eight (NS) cRNA and ccRNA with the samples used for sequencing (samples treated with oligo d(T)$_{25}$ beads). **e**, Percentage of segment eight cRNA missing the first A nucleotide in wild-type and T677A mutant virus samples. Data in panel **c, e** are presented as mean ± SD from three independent experiments. Statistical significance in **c** was determined by two-way Anova with Tukey's multiple comparisons test and in **e** by unpaired two-tailed t-test (ns=non-significant, **P*<0.05, *****P*<0.0001).

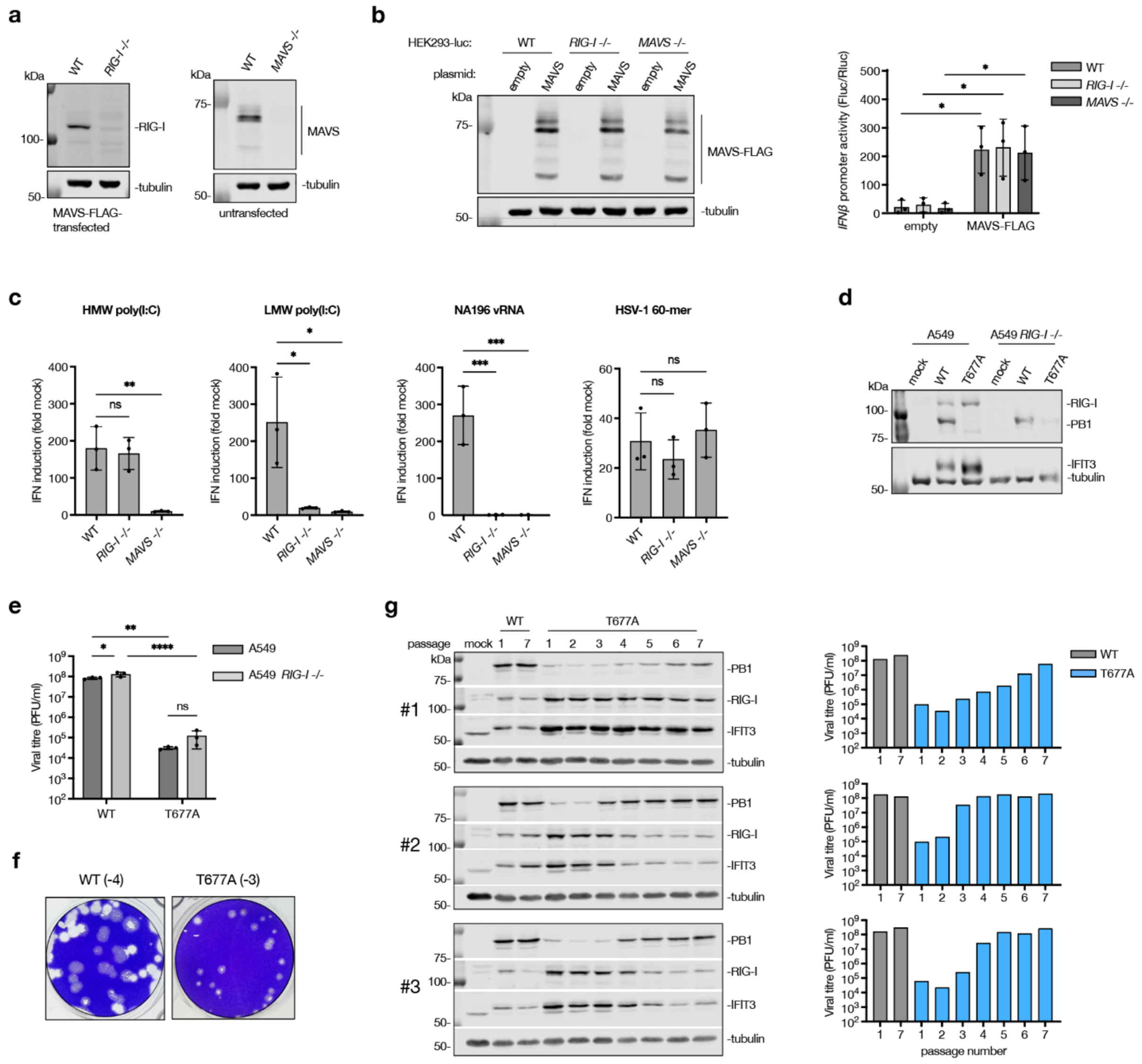


**Extended Data Fig. 4 *RIG-I* -/- and *MAVS* -/- HEK293-luc cell line characterization, T677A mutant virus infection and passaging in A549. a,** Efficiency of *RIG-I* and *MAVS* knockout was evaluated by western blotting. RIG-I expression in the left panel was induced by MAVS-FLAG overexpression for 24 hours. **b,** Wild-type, *RIG-I* -/-, and *MAVS* -/- HEK293-luc cells were transfected with either a MAVS-FLAG-expressing plasmid or an empty control plasmid, along with a *Renilla* luciferase reporter plasmid, for 24 hours. MAVS-FLAG expression was assessed by western blotting (left). The IFN-β luciferase reporter activity was expressed as a proportion of *Renilla* luciferase signal (right). **c**, Wild-type, *RIG-I* -/-, and *MAVS* -/- HEK293-luc cells were transfected with HMW poly(I:C), LMW poly(I:C), *in vitro* transcribed NA196 vRNA or HSV-1 60-mer DNA and IFN-β luciferase reporter activity was measured 24 hours post-transfection. **d**, A549 or A549 *RIG-I* -/- cells were infected with wild-type or T677A virus at an MOI of 0.01, or mock infected, for 48 hours. Protein expression was assessed by western blotting. A representative image from three independent experiments is shown. **e**, Viral titres in A549 or A549 *RIG-I* -/- cell culture supernatants collected from 48-hour infection at an MOI 0.01 were measured by plaque assay. **f**, Representative images from three independent experiments showing plaque sizes of viruses collected from infected HEK293-luc cells and grown on an MDCK monolayer. Sample dilutions are indicated. **g**, Wild-type and T677A mutant A/WSN/33 viruses were serially passaged seven times in A549 cells at an MOI of 0.01. Supernatants and protein samples were collected 48 hours post-infection for each passage. ISG and PB1 expression levels were assessed by western blotting (left) and viral titers were determined by plaque assay (right). Only passages 1 and 7 are shown for the wild-type virus. Images in panels **a** and **b** are representative of two replicates and in panel **d** and **f** of three replicates; in panel **g** all replicates are shown. Data in graphs are presented as mean ± SD from three independent experiments. Statistical significance was determined using two-way ANOVA with Šídák's multiple comparisons test (b) or Tukey's multiple comparisons test (e) and one-way ANOVA with Dunnett's multiple comparisons test (c) (ns=non-significant, *$P$<0.05, **$P$<0.01, ****$P$<0.0001).

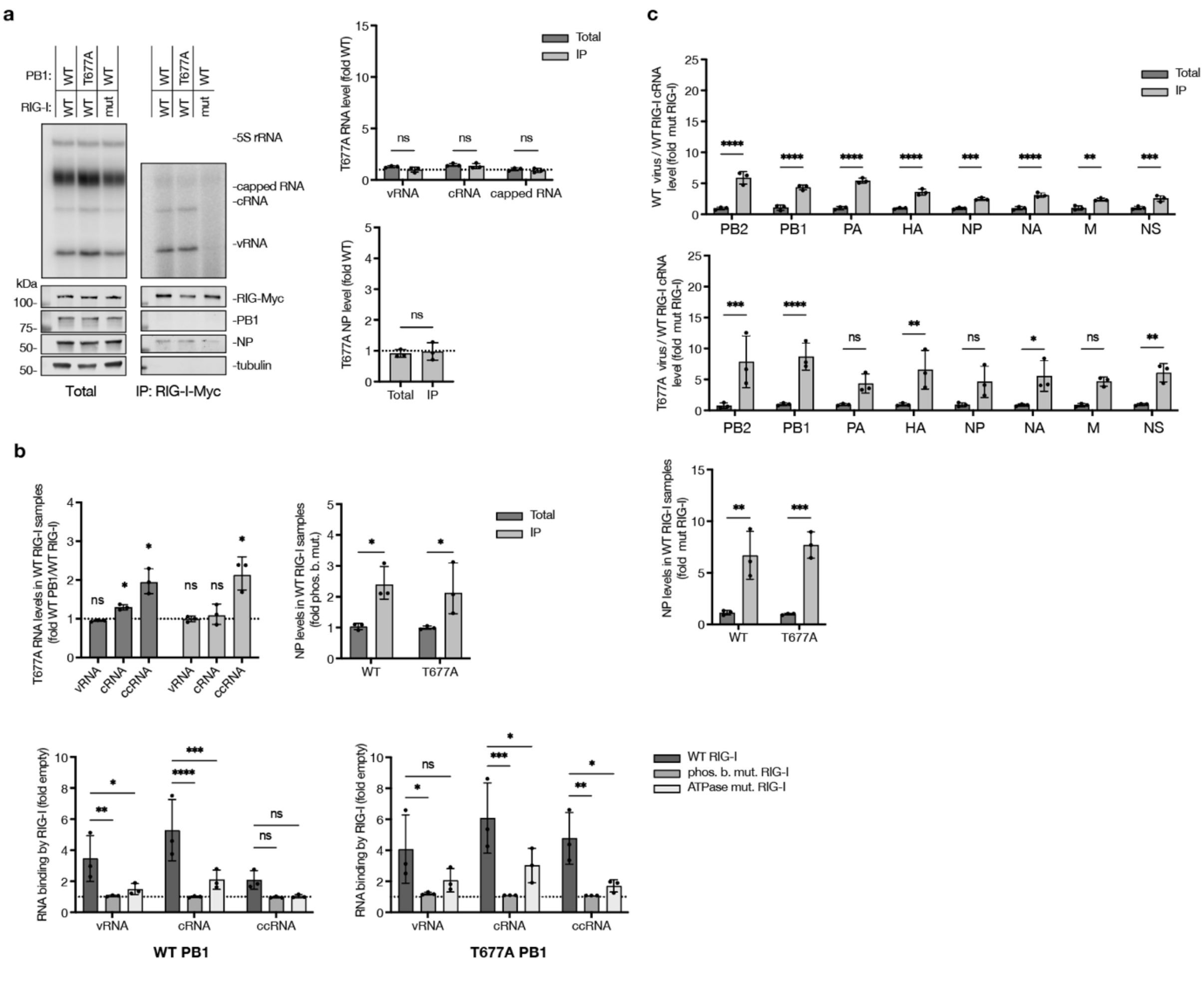

**Extended Data Fig. 5. RIG-I immunoprecipitation a**, Myc-tagged RIG-I was co-expressed with wild-type or T677A mutant RNP and segment 5 (NP), followed by immunoprecipitation. A phosphate-binding mutant of RIG-I was used as a negative control. RNA levels in total and immunoprecipitated (IP) fractions were measured by primer extension with internal +sense (NP 149+) and -sense (NP 149-) primers. Protein levels were assessed by western blotting. Shown are a representative image from three independent experiments. **b**, Quantification of RNA and nucleoprotein in total and IP fractions of RIG-I immunoprecipitation in RNP assay with NA196 template (top). Bottom panel shows quantifications of RNA levels in IP fraction in wild-type and mutant RIG-I conditions (shown as fold of empty plasmid levels). **c**, Quantification of cRNA (top two) and NP (bottom) levels in total and IP fractions of wild-type and T677A virus infected samples. All gel images are representative of three independent replicates. Data in all panels are shown as (mean ± SD) from three independent experiments. Statistical significance was determined using two-way ANOVA with Šídák's multiple comparisons test (**a**, top right; **b**, top right, **c**) or Dunnett's multiple comparisons test (**b**, bottom); one-sample t-test where each condition was compared to a theoretical mean of 1 (**b**, top left) and an unpaired two-tailed t-test (**a**, bottom right) (ns=non-significant, **P*<0.05, ***P*<0.01, ****P*<0.001, *****P*<0.0001).
